# Traip Mitotic Function Controls Brain Size

**DOI:** 10.1101/2021.07.07.451466

**Authors:** Ryan S. O’Neill, Nasser M. Rusan

## Abstract

Microcephaly is a developmental failure to achieve proper brain size and neuron number. Mutations in diverse genes are linked to microcephaly, including several with DNA damage repair (DDR) functions; however, it is not well understood how these DDR gene mutations limit brain size. One such gene is *TRAIP*, which has multiple known functions in DDR. We characterized the *Drosophila* ortholog *Traip*, finding that loss of *Traip* causes a brain-specific defect in the Mushroom Body (MB). *Traip* mutant (*traip*^-^) MBs had reduced size and fewer neurons, but no neurodegeneration, consistent with human primary microcephaly disorders. Reduced neuron numbers in *traip*^-^ were explained by premature caspase-dependent cell death of MB neuroblasts (MB-NBs). Many *traip*^-^ MB-NBs had prominent chromosome bridges in anaphase, along with polyploidy, aneuploidy, or micronuclei. We found no evidence for an interphase DNA repair role for Traip in MB-NBs; instead, proper MB development requires Traip function during mitosis, where Traip localizes to centrosomes and mitotic spindles. Our results suggest that proper brain size is ensured by the recently described role for TRAIP in unloading stalled replication forks in mitosis, which suppresses DNA bridges and neural stem cell death to promote proper neuron number. Further, the mitotic nature of *traip*^-^ MB-NB defects and Traip localization suggest a closer etiological link between DDR microcephaly genes like *Traip* and the centrosome/spindle-related genes more commonly associated with microcephaly.

## Introduction

Microcephaly is a developmental growth disorder characterized by reduced cerebral cortex size and neuron number. Mutations in about 40 genes are linked to heritable forms of microcephaly, ranging from MicroCephaly, Primary, Hereditary (MCPH; Jayaraman et al., 2018) where only brain size is reduced, to more severe forms like Seckel syndrome (SCKL) where both brain and body are reduced (Khetarpal et al., 2016). Most microcephaly disorders are linked to mutations in genes from one of two functional classes: centrosome and mitotic spindle functions, or DNA damage repair (DDR). Mutations in centrosome/mitotic spindle genes are thought to disrupt mitotic spindle structure, positioning, and orientation, leading to defects in chromosome segregation or cell cycle progression and ultimately neural progenitor cell (NPC) loss through premature differentiation or cell death (reviewed in Nano and Basto, 2017). Mutations in DDR genes are thought to increase DNA damage, activate checkpoint signaling and lead to increased genome instability, cell cycle lengthening, and the ultimate loss of NPCs (reviewed in Bianchi et al., 2018). However, in many cases microcephaly genes have not been directly studied in the context of brain development and the cellular etiology of microcephaly is inferred from studies in non-neuronal contexts. Thus, we aimed to study the microcephaly gene *TRAIP* in an animal to reveal the etiological mechanisms of disease.

*TRAIP* encodes a RING domain E3 ubiquitin ligase that localizes to the nucleus and functions in DDR. In humans, hypomorphic mutations in *TRAIP* cause SCKL with severe microcephaly and reduced body size (Harley et al., 2016). Cultured fibroblasts from *TRAIP* mutant patients have an elongated cell cycle and impaired DNA repair, suggesting that mutations in *TRAIP* cause microcephaly via reduced cell proliferation, premature differentiation, and increased apoptosis of NPCs (Harley et al., 2016). A mouse *TRAIP* null mutant is early embryonic lethal (Park et al., 2007) and *Drosophila* mutants for the *TRAIP* ortholog *nopo* (herein called *Traip*) are maternal effect lethal (homozygous null mothers lay eggs that arrest in the first few embryonic cell cycles; Merkle et al., 2009), whereas null mutant *C. elegans* are viable (Sonneville et al., 2019). No studies of *TRAIP* function in developing brains have been published.

TRAIP has several functions related to DNA repair (reviewed in Wu et al., 2020). TRAIP localizes to DNA double-strand breaks, where it recruits other factors and promotes H2B monoubiquitylation to stimulate repair (Han et al., 2019; Soo Lee et al., 2016). TRAIP also localizes to stressed replication forks (Feng et al., 2016; Hoffmann et al., 2016) where it removes DNA-protein crosslinks via ubiquitylation (Larsen et al., 2019) and regulates the choice between NEIL3 or FANC/BRCA-mediated repair pathways (Wu et al., 2019). In addition to these interphase DDR functions, TRAIP also promotes proper sister chromatid separation during mitosis. A cell entering mitosis may have under-replicated sister chromatids (URSCs) which contain loci that failed to complete replication during interphase, and these URSCs must be resolved to prevent DNA bridges during anaphase. TRAIP initiates the resolution of URSCs by ubiquitylating the MCM7 subunit of CMG helicase, thus triggering replication machinery unloading from stalled forks and allowing sister chromatid resolution via mitotic DNA repair synthesis (Deng et al., 2019; Priego Moreno et al., 2019; Sonneville et al., 2019; Villa et al., 2021).

However, most previous studies on TRAIP function were done in cell culture, *Xenopus* egg extracts, and the early embryos of *Drosophila* and *C. elegans*, and thus it is unclear which TRAIP functions are critical for neurogenesis. In this study we take advantage of the well characterized brain structure and genetic tractability of *Drosophila* to establish a model of *Traip* mutant microcephaly. In this context, Traip functions in NSCs during mitosis to suppress DNA bridges and prevent premature caspase-dependent cell death, pointing to a role in URSC resolution as critical for proper brain size.

## Results

### *Traip* is Required for Proper Mushroom Body Development

We generated a new full coding sequence deletion of *Traip* using CRISPR (*traip^Δ^*). We use *traip*^-^ to refer to *traip^Δ^* in combination with either the previously described null allele *traip^Exc142^* (Merkle et al., 2009) or the deficiency *Df(2R)Exel7153* (Parks et al., 2004), as both allelic combinations showed identical brain phenotypes.

To test whether *Traip* controls brain size, we used μ-computed tomography (μ-CT; Schoborg et al., 2019) to compare control and *traip*^-^ adult brains using normalized volumetric analysis (Figure 1A). While optic lobes were not different (Figure 1B), *traip*^-^ central brain volume was slightly reduced, and this reduction was rescued by a ubiquitously expressed GFP-tagged Traip transgene (*ubi-GFP::Traip*; Figure 1C). These initial results suggested that *Traip* is required for fully proper brain development.

**Figure 1.**
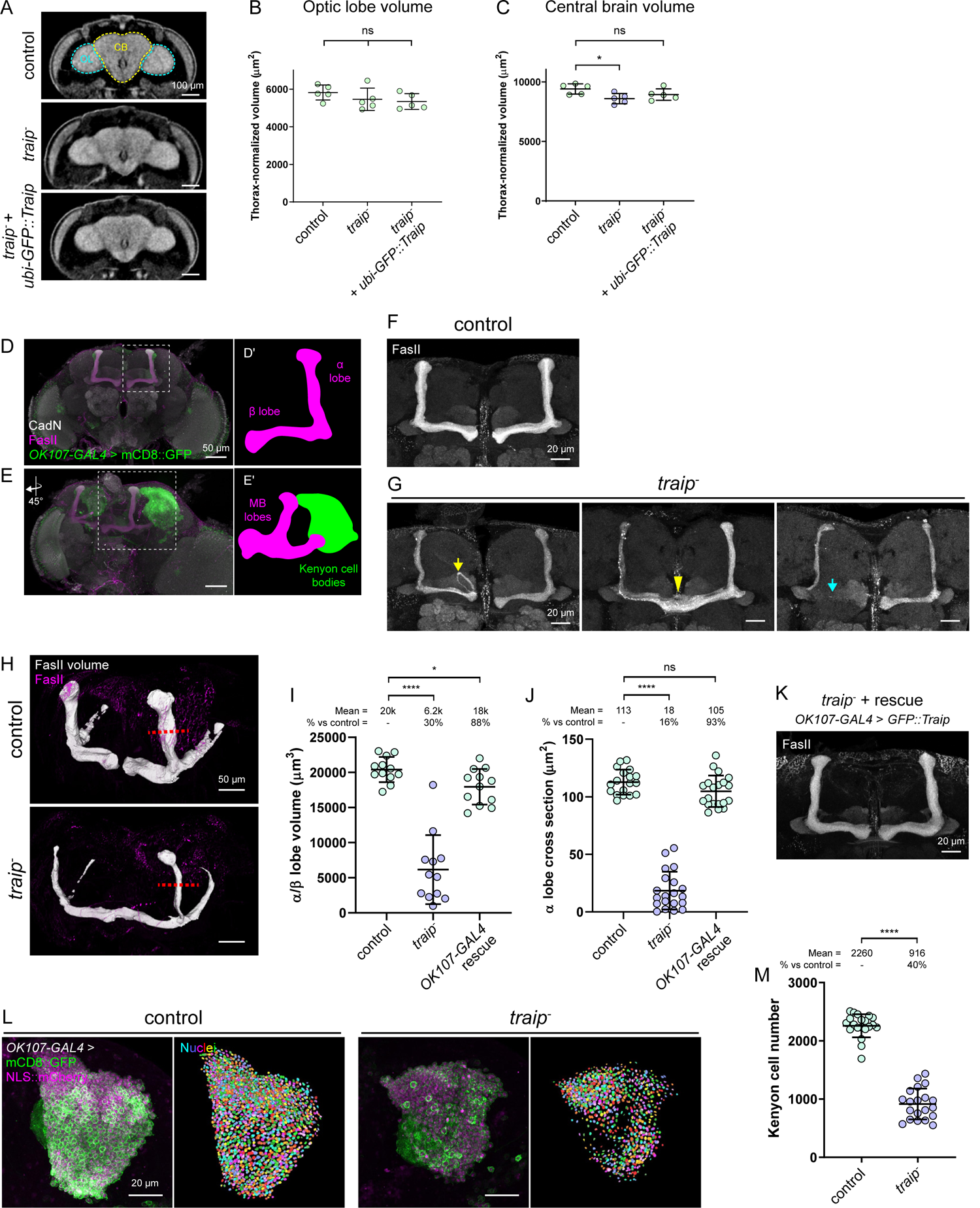
*Traip* is required for proper Mushroom Body structure. (A) μ-CT tomograms of control, *traip^-^*, and *traip^-^* + *ubi-GFP::Traip* adult brains. Optic lobes (OL) and central brain (CB) are highlighted on the control. (B) Thorax-normalized optic lobe volumes. N = 5 brains. (C) Thorax-normalized central brain volumes. N = 5 brains; * p = 0.025. (D) Maximum projection of adult brain stained for FasII (α/β lobes, magenta; cartoon in Dˈ) and CadN (neuropil, white) and labeled with *OK107-GAL4* > mCD8::GFP (green). (E) Oblique full brain volume projection stained for FasII, CadN, and labeled with *OK107-GAL4* > mCD8::GFP, showing KC bodies on the ventral posterior side projecting axons anteriorly to form the MB lobes (cartoon in Eˈ). (F) Control MBs stained for FasII with stereotypic L-shaped α/β lobes. (G) *traip^-^* MBs with reduced size, misguided axon tracts (yellow arrow, 50% *traip^-^* vs 0% control), midline fused MBs (yellow arrowhead, 93% *traip^-^* vs 14% control), and missing α or β lobes (cyan arrow, 29% *traip^-^* vs 0% control). (H) Control and *traip^-^* α/β lobe volumes (white) segmented from FasII staining (magenta). Red lines show position of cross-section measurements. (I) α/β lobe volume measurements. N ≥ 12 MBs. * p = 0.0145. (J) α lobe cross-section measurements. N ≥ 18 MBs. (K) MBs from *traip^-^* + *OK107-GAL4* > *GFP::Traip* rescue with wild-type morphology. (L) Machine learning segmentation of KCs. *OK107-GAL4* > mCD8::GFP (green) + NLS::mCherry (magenta, left panels) were used for KC nuclei segmentation and counting (multi-color, right panels) of controls and *traip^-^*. (M) KC numbers per hemisphere. KCs are reduced by 60% in *traip^-^*. N = 20 hemispheres. In all graphs, bars are mean ± standard deviation. Two-tailed t-test (B, C) and Mann-Whitney test (I, J, M) were used for significance. ns = not significant, **** p < 0.0001. Scale bars = 100 μm (A), 50 μm (D, E, H), 20 μm (F, G, K, L).

To determine if any specific central brain sub-region was disrupted, we examined neuropil (N-cadherin; CadN) and axon tracts (Neuroglian; Nrg) in *traip*^-^ brains. All brain regions appeared qualitatively similar (Figure S1A), except for a 100% penetrant size and structure defect in *traip*^-^ Mushroom Bodies (MBs), a pair of neuropils that mediate higher order functions in the insect brain (Figure 1D; Heisenberg, 2003; Modi et al., 2020). In *Drosophila*, each MB arises from four neuroblasts (MB-NBs), which divide continuously from embryonic development through late pupal stages to produce roughly 2500 Kenyon cells (KCs) per hemisphere (Figure 1E). These KCs are positioned on the posterior dorsal side of the brain and project axons in a stereotypic, bifurcated pattern to form the γ (first born), α’/β’, and α/β (last born) lobes of the MB (Figure 1D).

The MBs of *traip*^-^ adult brains were misshapen and reduced in size (Figure 1G), with additional axon-related defects: 50% with axon tracts exiting the lobes (0% in controls), 93% with lobes fused at the midline (14% in controls), and 29% with missing α or β lobes (0% in controls). *traip*^-^ α/β lobe volume, identified by FasII staining, was reduced by 70% (Figure 1H and 1I), and *traip*^-^ full MB volume, identified by mCD8::GFP expression driven in KCs via *OK107-GAL4*, was reduced by 37% (Figure S1B and S1C). The more severe size reduction of the last born, FasII-positive α/β lobes indicates that the *traip*^-^ MB defect progressively accumulates through development. Cross-sectional area was also reduced in FasII-positive α lobes by 84% (Figure 1J) and in *OK107-GAL4* > mCD8::GFP αˈ/α lobes by 54% (Figure S1D).

MB size reduction was consistent across all *Traip* null mutant combinations (Figure S1E), whereas hypomorphic mutants (Merkle et al., 2009) had wild-type MB size (Figure S1E and S1F). MB size was rescued by ubiquitously expressing GFP::Traip (Figure S1G and S1H), and importantly, by specifically expressing GFP::Traip in the MB-NBs and KCs via *OK107-GAL4* (Figure 1I-1K). Thus, MB size reduction is linked to cell-autonomous loss of Traip function in the MB-NBs and/or KCs.

Reduced *traip*^-^ MB lobe size suggests fewer axons and reduced KC number. We used machine learning to count KCs with *OK107-GAL4* > NLS::mCherry (Figure 1L). Each *traip*^-^ hemisphere contained an average of 916 KCs, a 60% reduction compared to controls with 2260 KCs (Figure 1M). Regression analyses showed a significant linear correlation between KC number and both MB volumes (Figure S2I) and cross sectional areas (Figure S2J). Having shown that α lobe cross-sectional area measurement was a suitable proxy for KC number, we used this measurement for the remainder of this study.

### *Traip* is Required in Mushroom Body Neuroblasts During Development

To determine whether the *traip*^-^ MB size reduction was developmental (primary microcephaly-like) or neurodegenerative (secondary microcephaly-like; Passemard et al., 2013), we investigated pre-adult stages (Figure 2A). MB size was not significantly reduced in *traip*^-^ larvae, but was markedly reduced through pupal stages (Figure 2B). Thus, *traip*^-^ is primary microcephaly-like, as MB size reduction arises during development.

**Figure 2.**
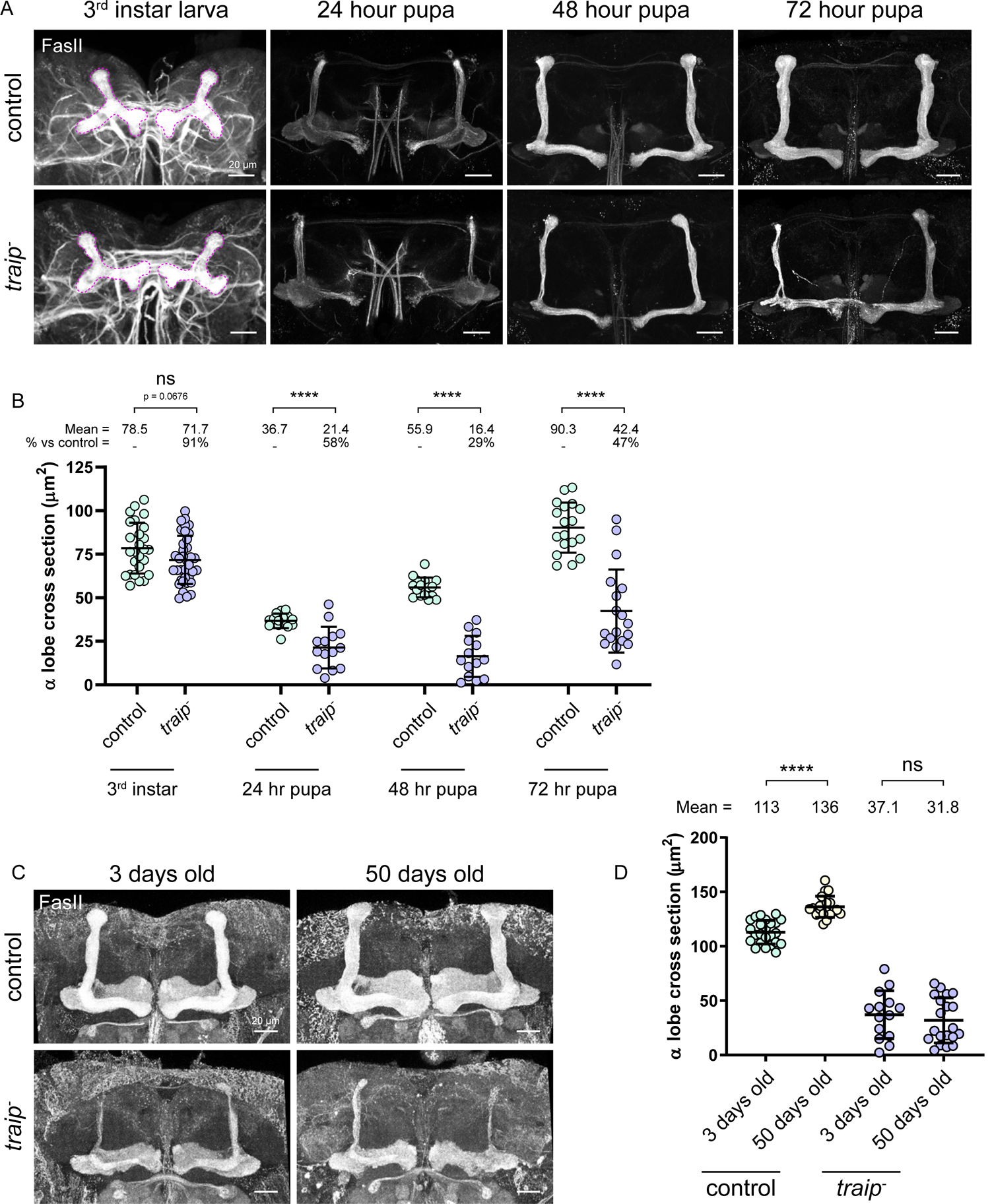
*Traip* is required for MB development. (A) Control (top row) and *traip^-^* (bottom row) MBs from (left to right) 3^rd^ instar larvae, 24 hours APF, 48 hours APF, and 72 hours APF pupae stained for FasII. 3^rd^ instar larval MBs are highlighted in magenta; note that in larval stage, FasII labels the γ lobes, and α/β lobes are yet to be born. (B) MB lobe cross-section measurements of control and *traip^-^* developmental stages. 3^rd^ instar larval MBs of *traip^-^* are not significantly reduced compared to controls. In all pupal stages, *traip^-^* are significantly reduced compared to controls. Note that 24 hours APF MBs appear smaller than larval MBs in part due to extensive morphological remodelling in early metamorphosis (Lee et al., 1999). N ≥ 14 MBs. (C) Control (top row) and *traip^-^* (bottom row) MBs from 3 day old adults (left column) and 50 day old adults (right column) stained for FasII. (D) α lobe cross-section measurements show that control MBs increase in size with age, whereas *traip^-^* MBs do not change. N ≥ 14 MBs. Two-tailed t-test was used for significance. ns = not significant, **** p < 0.0001. Scale bars = 20 μm.

To investigate a possible role in neurodegeneration we compared the MBs of three day old and 50 day old adults (Figure 2C). MB size increased slightly with age in controls (Figure 2D), reflecting neuropil reorganization in response to life experience (Heisenberg et al., 1995). In contrast, *traip*^-^ MBs neither grew nor shrunk with age (Figure 2D). Thus, *traip*^-^ MB size reduction is not neurodegenerative and does not reflect secondary microcephaly.

We hypothesized that *traip*^-^ reduced KC number and MB size could arise in two ways: 1) a defect in the highly mitotic MB-NBs resulting in fewer KCs born, and 2) a defect in the differentiated KCs resulting in cell death. We began by exploring which cell types express Traip, using CRISPR to knock-in *mNeonGreen* at the endogenous *Traip* locus (*mNG::Traip*), which resulted in a fly line with wild-type MB morphology (Figure S2A). mNG::Traip was expressed in all 3^rd^ instar larval central brain NBs (CB-NBs), which include the MB-NBs (Figure 3A). In addition, mNG::Traip expression persisted in the mitotic ganglion mother cells (GMCs) and their immediate daughter neurons (Figure 3B). However, mNG::Traip was absent from regions of mature neurons in larval brains (Figure 3A), and was not detectable in adult brain neurons (Figure S2B). These data suggest that Traip primarily functions in proliferating brain cells.

**Figure 3.**
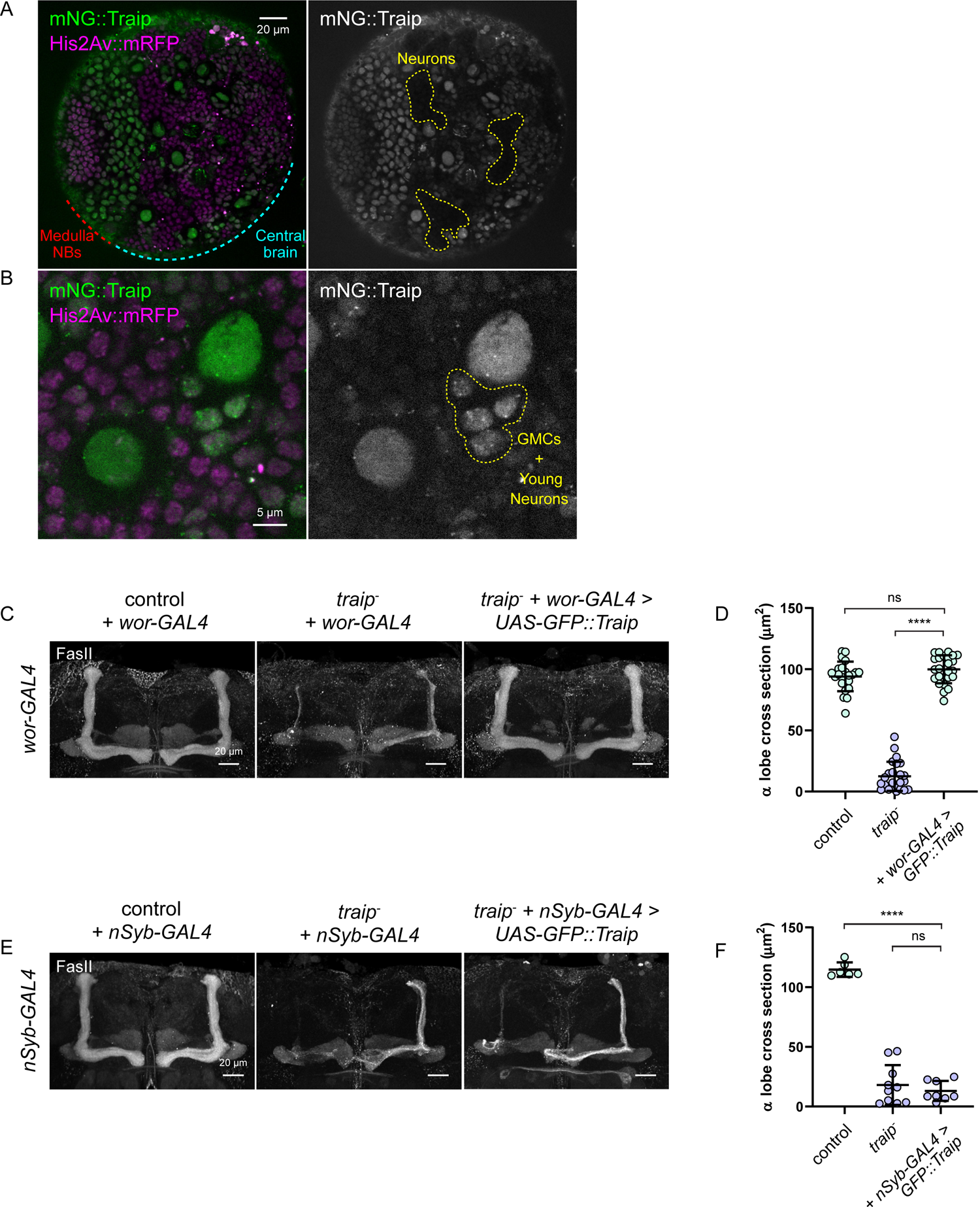
*Traip* is required in neuroblasts. (A) Endogenously tagged mNG::Traip (green, gray) and His2Av::mRFP (magenta) in the 3^rd^ instar larval brain. mNG::Traip is highly expressed in the proliferating cells of the medulla (red zone) and central brain regions (cyan zone), but is absent from areas dominated by neurons (yellow outlines). Cell types were determined by position and morphology as mNG::Traip fluorescence does not survive fixation. (B) High resolution imaging of mNG::Traip localized to the nuclei of interphase CB-NBs, and persisting in daughter GMCs and young neurons (yellow outline). (C) MBs from control, *traip^-^*, and *UAS-GFP::Traip* rescue, all with *wor-GAL4*, stained for FasII. (D) α lobe cross-section measurements show that *traip^-^* + *wor-GAL4* > *GFP::Traip* rescue have wild-type MB size. N ≥ 22 MBs. (E) MBs from control, *traip^-^*, and *UAS-GFP::Traip* rescue, all with *nSyb-GAL4*, stained for FasII. (F) α lobe cross-section measurements show that *traip^-^* + *nSyb-GAL4* > *GFP::Traip* have reduced MB size. N ≥ 6 MBs. Two-tailed t-test was used for significance. ns = not significant, **** p < 0.0001. Scale bars = 20 μm (A, C, E), 5 μm (B).

To determine which cell types require Traip function, we used cell type-specific expression of GFP::Traip in attempt to rescue *traip*^-^ MB size. First, we used *wor-GAL4* to drive *UAS-GFP::Traip* in NBs, which rescued MB size (Figure 3C and 3D). However, this did not fully exclude a role for Traip in neurons, as *wor-GAL4* > GFP::Traip was also found in the GMCs and newly born neurons (Figure S2C). We next tried expressing GFP::Traip in post-mitotic neurons using *elav-GAL4*, but discovered strong expression in the NBs and GMCs (Figure S2D). Finally, we used *nSyb-GAL4* to specifically drive GFP::Traip expression in mature neurons (Figure S2E), which resulted in failure to rescue MB size (Figure 3E and 3F). Thus, Traip function in neurons alone is insufficient to prevent microcephaly. We conclude that the primary function for Traip is in proliferating brain cells (NBs and GMCs), albeit we could not fully rule out the possibility that undetectable levels of Traip also function in post-mitotic neurons.

### *Traip* Mutant Mushroom Body Neuroblasts are Lost via Caspase-Dependent Cell Death

We next hypothesized that *traip*^-^ reduction in KC number and MB size is due to premature loss of MB-NBs. The KCs of each MB arise from four MB-NBs that are easily identifiable at 24 hours after pupal formation (APF), as nearly all other NBs are lost by this stage (Ito and Hotta, 1992; Truman and Bate, 1988). Controls maintained four MB-NBs throughout pupal development (Figure 4A and 4B). In contrast, *traip*^-^ progressively lost their MB-NBs, with an average 2.2 MB-NBs per hemisphere at 24 hours APF decreasing to 0.9 by 72 hours APF, and this loss was rescued by *OK107-GAL4* > *UAS-GFP::Traip* (Figure 4A and 4B).

**Figure 4.**
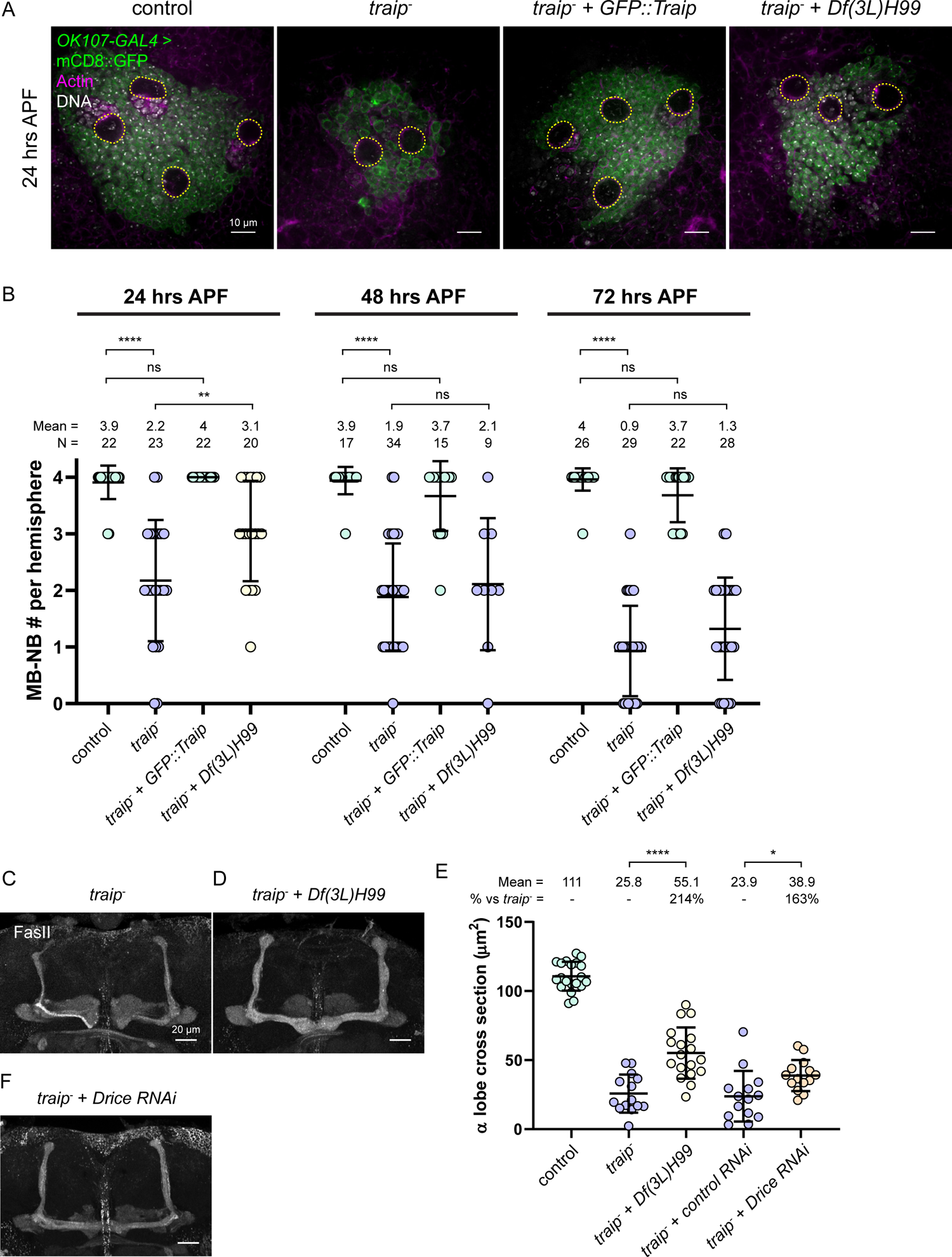
*Traip* suppresses MB-NB cell death. (A) Fields of KCs and MB-NBs labelled with *OK107-GAL4* > mCD8::GFP (green), phalloidin (magenta), and DAPI (gray) from (left to right) control, *traip^-^*, *traip^-^* + *OK107-GAL4* > *GFP::Traip* rescue, and *traip^-^* + *Df(3L)H99* 24 hours APF pupae. NBs are highlighted in yellow. Scale bars = 10 μm. (B) MB-NB number per brain hemisphere for control, *traip^-^*, *traip^-^* + *GFP::Traip* rescue, and *traip^-^* + *Df(3L)H99* at 24, 48, and 72 hours APF pupal stages. Two-tailed Mann-Whitney test was used to determine significance. ns = not significant, ** p = 0.0069, **** p < 0.0001. N is reported above each column. (C, D, F) Maximum projections of adult MBs stained for FasII. Genotypes are: *traip^-^* (C); *traip^-^* + *Df(3L)H99* (D); *traip^-^* + *OK107-GAL4* > *Drice* RNAi (F). Scale bars = 20 μm. (E) α lobe cross-section measurements of control, *traip^-^*, *traip^-^* + *Df(3L)H99*, *traip^-^* + control RNAi and *traip^-^* + *Drice* RNAi. Two-tailed t-test was used for significance. * p = 0.0149, **** p < 0.0001. N ≥ 14 MBs.

To explore the role of caspase-dependent cell death in MB-NB loss we used *Df(3L)H99*, which deletes four pro-apoptotic genes (*grim*, *rpr*, *hid*, and *skl*; Abbott and Lengyel, 1991). MB-NB loss was suppressed in *traip*^-^ with *Df(3L)H99* compared to *traip*^-^ alone at 24 hours APF (3.1 vs 2.2; Figure 4A and 4B), although the effect is not observed in later stages. *Df(3L)H99* also improved *traip*^-^ MB size more than 2-fold (Figure 4D and 4E). Since *Df(3L)H99* deletes other genes, we also used RNAi against the effector caspase *Drice*, which improved *traip*^-^ MB size by 1.6-fold (Figure 4E and 4F). Together, these data support a model where Traip prevents premature caspase-dependent cell death of MB-NBs to ensure proper KC number and MB morphology.

### *Traip* Suppresses Chromosome Bridges During Anaphase

After noticing abnormal nuclei while scoring *traip*^-^ MB-NB number, we characterized nuclear phenotypes in 24 hours APF MB-NBs to gain insight into defects potentially upstream of MB-NB death. Control MB-NBs had nuclei with relatively smooth, spherical nuclear lamina morphology and no other defects (100%; Figure 5A). In contrast, some *traip*^-^ MB-NBs appeared apoptotic, with abnormally condensed DAPI staining (11%; Figure 5B), while others frequently had nuclear defects including ruffled, convoluted nuclear lamina morphology (12%; Figure 5C), micronuclei (11%; Figure 5D), multiple nuclei (9%; Figure 5E), or extremely large nuclei suggestive of polyploidy (9%; Figure 5F). These *traip*^-^ phenotypes were rescued by *OK107-GAL4* > *UAS-GFP::Traip* (Figure 5G).

**Figure 5.**
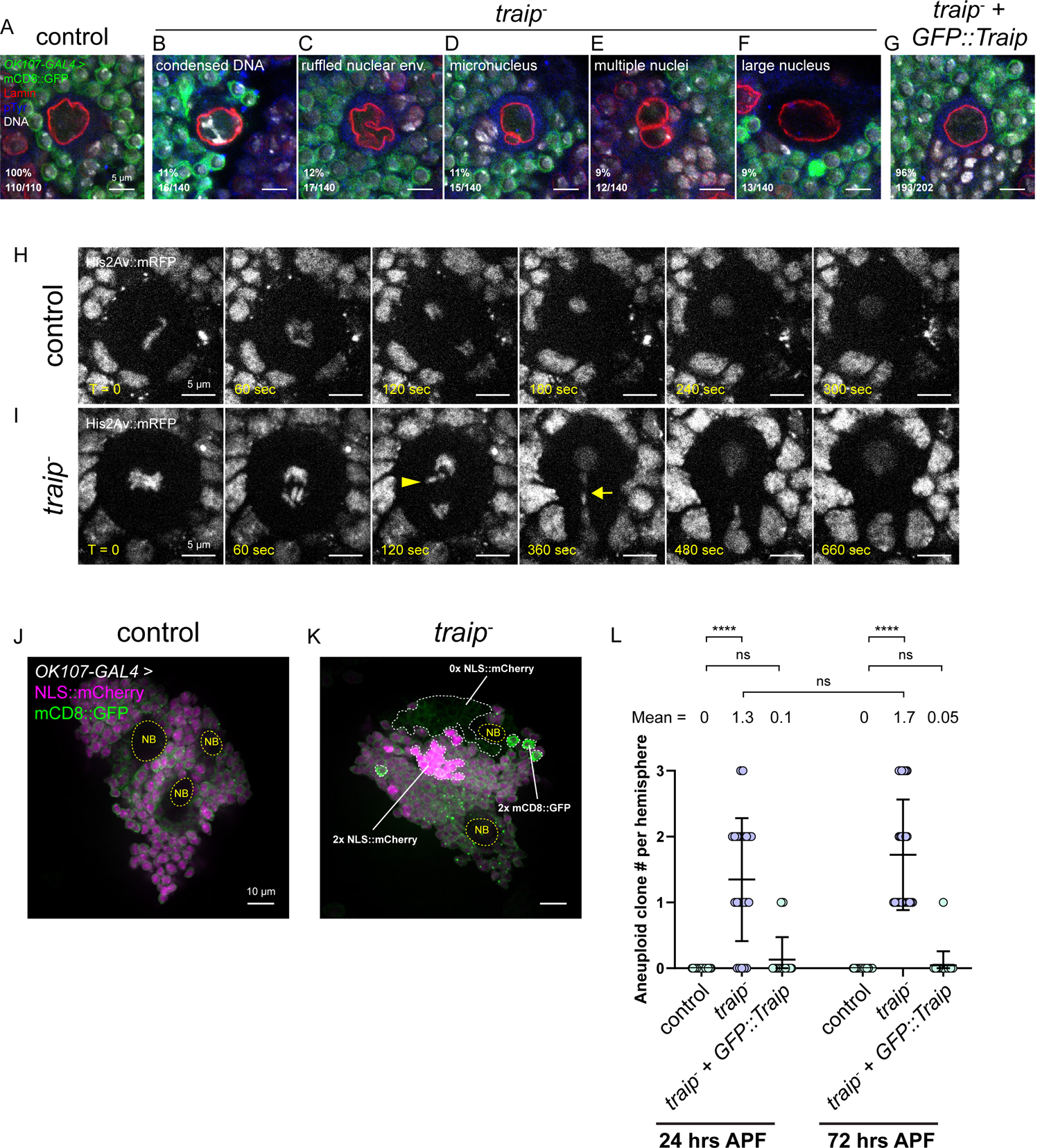
*Traip* suppresses multinuclear phenotypes and mitotic DNA bridges. (A-F) MB-NBs from 24 hours APF pupal brains labelled with *OK107-GAL4* > mCD8::GFP (green) and stained for Lamin (red), DNA (gray), and phospho-Tyrosine (blue). (A) 100% (110/110) of control MB-NBs have relatively smooth, spherical nuclear laminar morphology and no additional nuclear defects. (B) 11% (16/140) of *traip^-^* 24 hours APF MB-NBs appeared apoptotic, with abnormally condensed DAPI staining. (C) 12% (17/140) of *traip^-^* MB-NBs have irregular, ruffled nuclear envelope morphology. (E) 11% (15/140) of *traip^-^* MB-NBs have micronuclei. (E) 9% (12/140) of *traip^-^* MB-NBs have multiple nuclei. (F) 9% (13/140) of *traip^-^* MB-NBs appeared polyploid, with abnormally large nuclei. (G) Most MB-NBs from *traip^-^* + *OK107-GAL4* > *GFP::Traip* rescues have wild-type nuclear envelope morphology (96%, 193/202), and 4% (9/202) had ruffled nuclear envelope morphology. (H) Control 3^rd^ instar larval CB-NB expressing His2Av::mRFP. Chromosomes segregate effectively during mitosis. N = 10 NBs. See Movie 1. (I) 26% (5/19) of *traip^-^* CB-NBs form chromosome bridges during anaphase. A yellow arrowhead points to a chromosome as it pulls away in mid-anaphase, and a yellow arrow points to the chromosome bridge stretched across the constricting furrow. See Movie 2. (J) Control 24 hours APF pupae expressing *OK107-GAL4* > NLS::mCherry + mCD8::GFP show uniform fluorescence levels of both markers across the field of KCs, except for NBs (yellow outlines) and their immediate daughters which have reduced fluorescence. (K) *traip^-^* pupae contain patches of presumably aneuploid KCs with either double or no fluorescence for one or both markers (white outlines). (L) Number of aneuploid KC clones per brain hemisphere in control, *traip^-^*, and *traip^-^* + *OK107-GAL4* > *GFP::Traip* rescue at 24 and 72 hours APF. Two-tailed t-test was used for significance. ns = not significant, **** p < 0.0001. N ≥ 22 hemispheres. Scale bars = 5 μm (A-I), 10 μm (J, K).

As these nuclear defects could arise during mitosis, we used high resolution live fluorescence microscopy to analyze mitotic dynamics of 3^rd^ instar larval CB-NBs. Unlike control NBs (Figure 5H and Movie 1), 26% of *traip*^-^ NBs had prominent chromosome bridges, where sister chromatids appeared attached and did not effectively segregate, but rather became stretched across the midzone through anaphase before eventually separating (Figure 5I, Movie 2). Further, *traip*^-^ larval brains often had cells with abnormally large nuclei (Figure S3A), reminiscent of the suspected polyploid cells seen in *traip*^-^ pupal brains (Figure 5E). We reasoned that *traip*^-^ mitotic DNA bridges could block cytokinesis, resulting in multiple nuclei or polyploidy, or else lead to chromosome breakage, resulting in micronuclei.

We also found evidence suggesting persistent aneuploidy in *traip*^-^ MB-NBs. Control pupae expressing both mCD8::GFP and NLS::mCherry via *OK107-GAL4* had similar fluorescence levels across the field of mature KCs (Figure 5J). In contrast, *traip*^-^ pupae often contained one or more clusters of KCs that had either double the signal or no signal for the GFP or mCherry marker (Figure 5K). These aberrant KC clusters were reminiscent of clonal cell clusters, suggesting that their MB-NB progenitors became aneuploid, either losing or gaining a copy of one transgene, and then continued generating daughter cells. These clones were never observed in controls, whereas *traip*^-^ brains had an average of 1.3 clones per hemisphere at 24 hours and 1.7 clones per hemisphere at 72 hours APF (Figure 5L). The average number of KCs per clone in *traip*^-^ brains increased from 15 at 24 hours to 37 at 72 hours APF (Figure S3B), with some clones containing hundreds of KCs. *OK107-GAL4* > *UAS-GFP::Traip* suppressed clone formation (Figure 5L). These data are consistent with *traip*^-^ mitotic DNA bridges leading to aneuploid MB-NBs which often continue producing daughters before eventually being lost.

We next used live imaging to explore whether *traip*^-^ NBs were defective in mitotic timing, but found no difference in the duration from prophase onset to completion of furrow constriction for CB-NBs (Figure S3C). Additionally, the presence or absence of DNA bridges in *traip*^-^ NBs did not affect mitotic timing from anaphase onset to furrow constriction (Figure S3D), although we could not rule out a possible delay or failure in abscission. In fixed larval brains, the mitotic index of control and *traip*^-^ CB-NBs were similar (Figure S3E). In 24 hours APF brains, the mitotic index of *traip*^-^ MB-NBs was lower than controls (20% *traip*^-^ vs 35% controls); however, when MB-NBs with nuclear defects (Figure 5B-F) were excluded from this analysis, the mitotic index of phenotypically normal *traip^-^* MB-NBs was not different from controls (32% *traip^-^* vs 35% controls; Figure S3F). Thus, the reduced mitotic index of *traip*^-^ MB-NBs could be explained by an inability of polyploid, multinucleate, and apoptotic cells to enter mitosis rather than a function for Traip in directly controlling mitotic or cell cycle timing.

### *Traip* Functions to Resolve Under-Replicated Sister Chromatids

TRAIP has a known role in resolving under-replicated sister chromatids (URSCs) at mitotic onset (Deng et al., 2019; Priego Moreno et al., 2019; Sonneville et al., 2019), and URSCs are predicted to form a special class of mitotic DNA bridges called ultrafine DNA bridges (UFBs; Liu et al., 2014). To test for UFBs we generated transgenes encoding fluorescently tagged FancD2; human FANCD2 localizes as puncta at the points where UFBs connect with sister chromatids (Naim and Rosselli, 2009), and although FancD2 localization has not been well-characterized in *Drosophila*, there is precedence for invisible DNA tethers in *Drosophila* cells (Royou et al., 2010). We found that mNG::FancD2 puncta localized to mitotic DNA bridges in many *traip*^-^ NBs (7/12; Figure 6B), in addition to weakly localizing to mitotic chromosomes in both controls and *traip*^-^ NBs (Figure 6A, 6B). The observed FancD2 on chromosome bridges suggests the presence of UFBs and is consistent with a role for Traip in resolving URSCs at mitotic onset.

**Figure 6.**
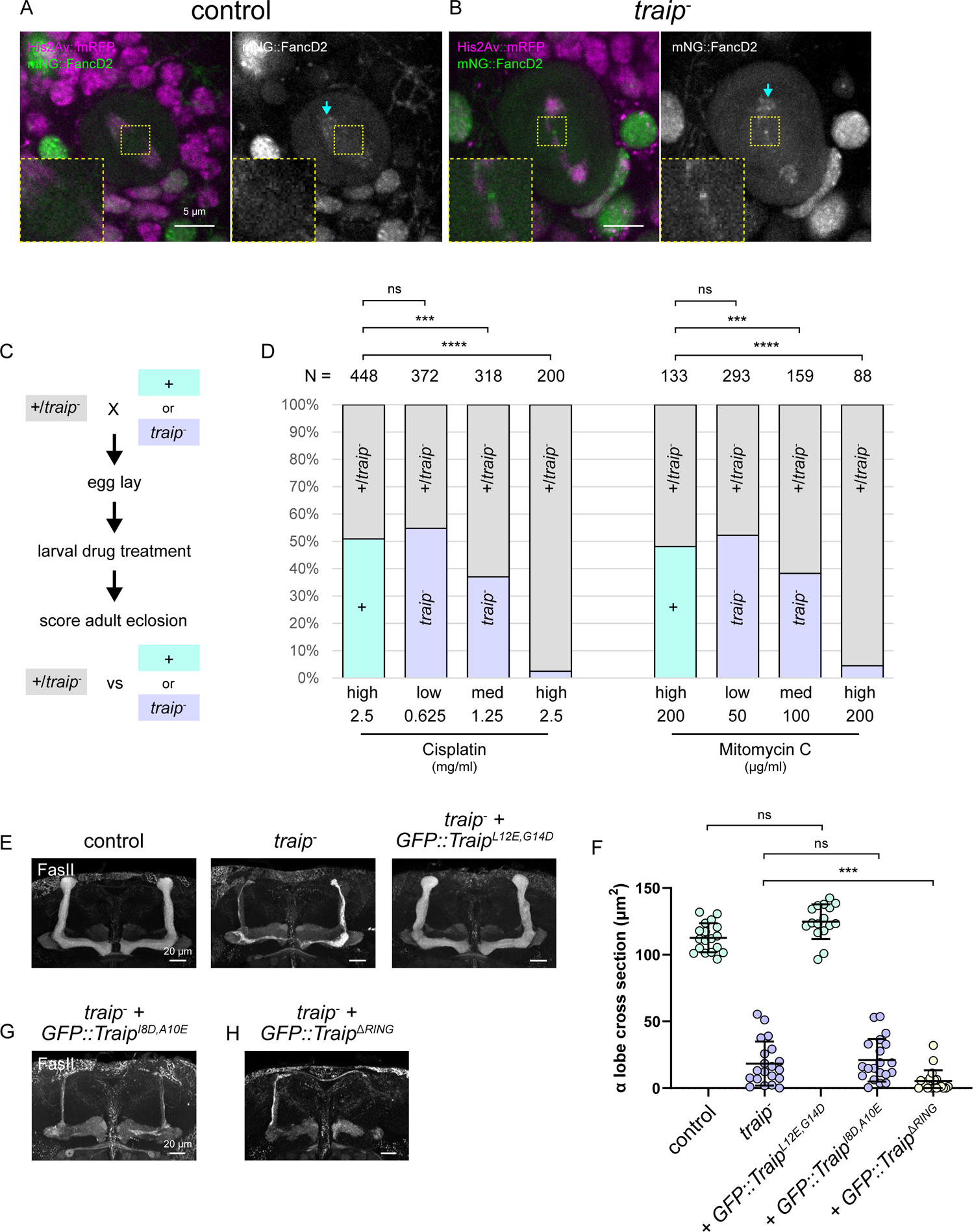
*Traip* is required for inter-strand crosslink repair. (A, B) 3^rd^ instar larval CB-NBs expressing mNG::FancD2 (green, gray) with His2Av::mRFP (magenta). 7/12 *traip^-^* NBs with mitotic DNA bridges had mNG::FancD2 puncta localized on the bridge (B, inset). Both controls and *traip^-^* have weak localization of mNG::FancD2 on mitotic chromosomes (cyan arrows). Scale bars = 5 μm. (C) Experimental setup for drug treatment survival assays. *+/ traip^-^* (*traip^Exc142^/CyO*) females were mated to either *+* (cyan) or *traip^Δ^* (purple) males. The females laid eggs for 24 hours, which were aged for another 24 hours before adding drug. The larvae developed and adult eclosion was scored. Offspring were either *+/ traip^-^ (+/traip^Δ^*) vs + (*+/CyO*) for control crosses (cyan), or *+/ traip^-^* (*traip^Δ^/CyO*) vs *traip^-^* (*traip^Δ^/traip^Exc142^*) for *traip^-^* crosses (purple). (D) Drug treatment survival assay results. Control crosses produced roughly similar numbers of *+* vs *+/ traip^-^* offspring with high doses of inter-strand crosslinking agents Cisplatin or Mitomycin C. *traip^-^* crosses also produced similar numbers of *traip^-^* vs *+/ traip^-^* offspring with low doses of either drug, but medium doses were semi-lethal and high doses were almost fully lethal to *traip^-^* offspring. Chi-squared tests were used to determine significance. ns = not significant, *** p < 0.001, **** p < 0.0001. (E, G, H) MBs from stained for FasII. Genotypes are: control, *traip^-^*, *traip^-^* + *GFP::Traip^L12E,G14D^* (E); *traip^-^* + *GFP::Traip^I8D,A10E^* (G); *traip^-^* + *GFP::Traip^ΔRING^* (H). Transgene expression was driven by *OK107-GAL4*. Scale bars = 20 μm. (F) α lobe cross-section measurements of control, *traip^-^*, and *traip^-^* + *GFP::Traip* RING mutant variants. Two-tailed t-tests were used for significance. *** p = 0.0004. N ≥ 16 MBs/genotype.

To better understand the mechanism of URSC and mitotic DNA bridge formation, we used drug treatments to induce replication stress via various mechanisms in control and *traip*^-^ larvae and assayed their survival to adult (Figure 6C). A major source of URSCs is late replicating regions, where replication forks fail to converge before mitotic entry (Liu et al., 2014); hydroxyurea increases the number of late replicating regions by depleting the nucleotide pool and inhibiting replication (Bianchi et al., 1986). Surprisingly, *traip*^-^ mutants were not sensitive to hydroxyurea treatment (Figure S4A), suggesting that late replicating region-induced URSCs are not the major target for Traip.

Another source of URSCs is inter-strand crosslinks, which block replication machinery and cause forks to stall; both Cisplatin and Mitomycin C are inter-strand crosslinking agents (Deans and West, 2011). Whereas controls tolerated high doses of Cisplatin and Mitomycin C, *traip*^-^ animals only tolerated low doses, with higher doses causing lethality (Figure 6D), likely due to defects in imaginal disc proliferation (Figure S4B). The MBs of *traip^-^* adults raised on medium doses of Cisplatin or Mitomycin C were not affected (Figure S4C, S4D); however, this may be due to a failure of these drugs to cross the blood brain barrier, as is the case in mammals (Gregg et al., 1992; Reddy and Randerath, 1987). Nonetheless, these experiments indicate that Traip functions to resolve DNA crosslink-induced damage.

TRAIP triggers replication machinery unloading via ubiquitylation of MCM7 (Wu et al., 2019), and therefore requires an intact RING domain for E2 conjugating enzyme binding. Thus, we tested whether the RING domain is required for Traip function in MB development. *GFP::Traip^L12E,G14D^*, which is a control mutant predicted to have a functional RING domain, fully rescued *traip*^-^ MB size (Figure 6E and 6F). In contrast, *GFP::Traip^I8D,A10E^*, which is predicted to have a non-functional RING-E2 binding interface, failed to rescue *traip*^-^ MB size (Figure 6F and 6G). Interestingly, *GFP::Traip^ΔRING^* enhanced the *traip*^-^ MB size defect (Figure 6H), indicating a dominant negative effect of deleting the RING domain. These results show that *Traip* function in MB-NBs is RING-dependent, consistent with an E3 ligase function and role in replication machinery unloading.

### Proper MB Development requires mitotic but not interphase Traip function

We discovered a novel mitotic localization for Traip while characterizing mNG::Traip expression. In interphase, mNG::Traip localized to the nucleus, as previously described (Feng et al., 2016; Harley et al., 2016; Soo Lee et al., 2016). However, in mitosis mNG::Traip localized to mitotic spindles and concentrates at the centrosomes (Figure 7A and Movie 3). High resolution imaging revealed that mNG::Traip forms small puncta that travel poleward along the spindles (Figure 7B and Movie 4) and also coalesce at the cytokinetic furrow and midbody (Figure 7C) in late mitosis.

**Figure 7.**
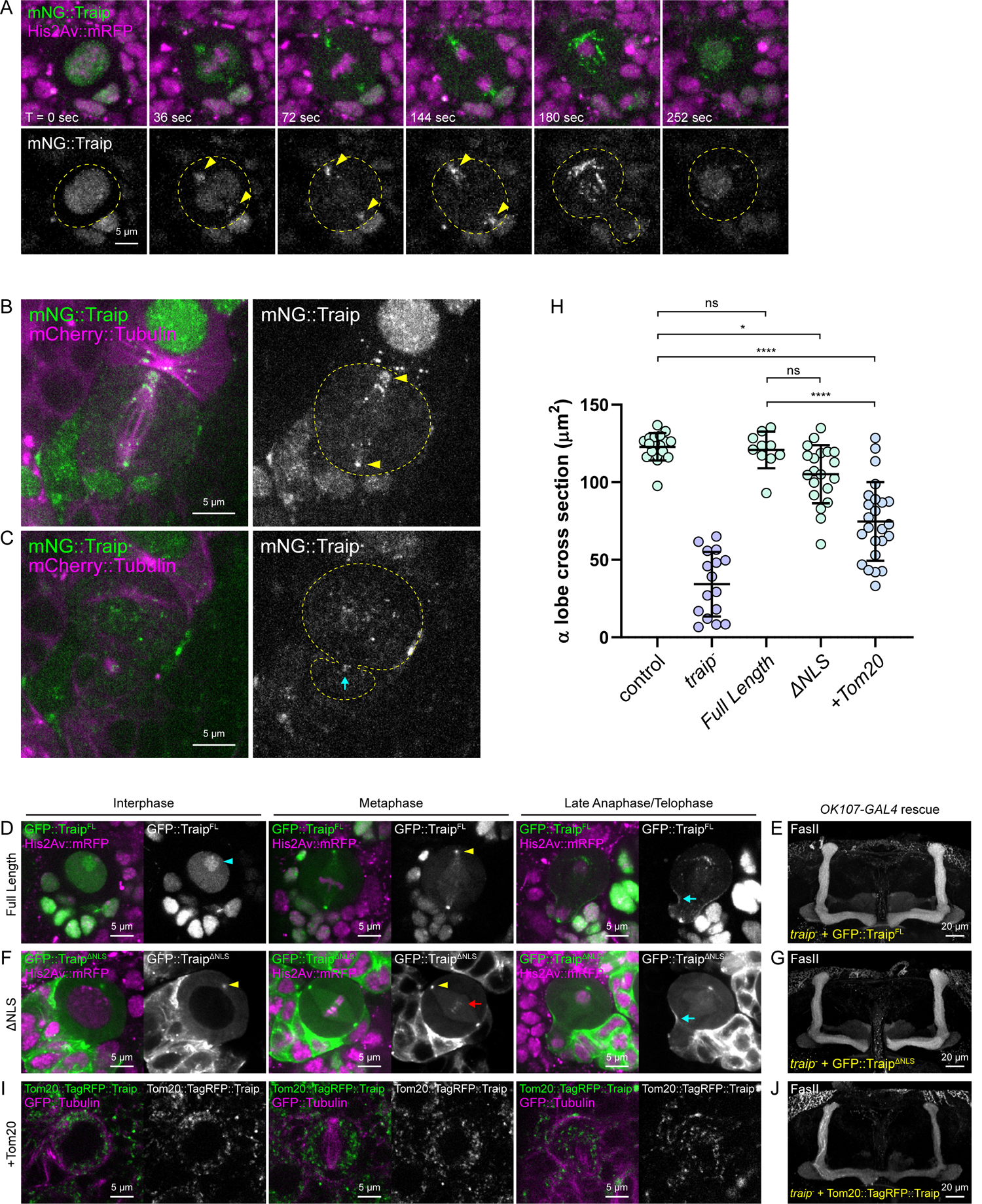
Traip has dynamic localization in mitosis. (A) Live imaging of mNG::Traip (green, gray) with His2Av::mRFP (magenta) in 3^rd^ instar larval CB-NB mitosis. At mitotic onset (36 sec) mNG::Traip is released from the nucleus and localizes to the centrosomes (yellow arrowheads) and spindle. See Movie 3. (B-C) High magnification live imaging of mNG::Traip (green, gray) with *wor-GAL4* > mCherry::Tubulin (magenta). mNG::Traip forms puncta that travel poleward on the spindle and coalesce at the centrosomes (yellow arrowheads, B), and coalesces at the cytokinetic furrow and midbody in late mitosis (cyan arrow, C). See Movie 4. (D, F) Localization of GFP::Traip variants (green) expressed via *wor-GAL4* with His2Av::mRFP (magenta) in CB-NBs during interphase, metaphase, and late mitosis. (D) GFP::Traip^Full Length^ has nucleolar localization in interphase (cyan arrowhead), centrosome localization in mitosis (yellow arrowhead), and furrow localization in late mitosis (cyan arrow). (F) GFP::Traip^ΔNLS^ lacks nuclear localization in interphase, but has centrosome localization in interphase and mitosis (yellow arrowhead), furrow localization in late mitosis (cyan arrow), and also localizes to metaphase chromosomes (red arrow). (E, G, J) MBs stained for FasII. Genotypes are: *traip^-^* + *GFP::Traip^FL^* (E); *traip^-^* + *GFP::Traip^ΔNLS^* (G); and *traip^-^* + *Tom20::TagRFP::Traip* (J). Transgene expression was driven by *OK107-GAL4*. (H) α lobe cross-section measurements of control, *traip^-^*, *traip^-^* + *GFP::Traip^FL^*, *traip^-^* + *GFP::Traip^ΔNLS^*, and *traip^-^* + *Tom20::TagRFP::Traip*. Ordinary one-way ANOVA with Tukey’s test was used for significance. ns = not significant, * p = 0.0448, **** p < 0.0001. N ≥ 16 MBs. (I) Localization of Tom20::TagRFP::Traip (green) with GFP::Tubulin (magenta) in CB-NBs. Tom20::TagRFP fails to localize to the nucleus in interphase, and is absent from the centrosome and spindle region in mitosis. Scale bars = 5 μm (A-C, D, F, I), 20 μm (E, G, J).

To characterize regions within Traip responsible for its mitotic localizations, we expressed GFP::Traip variants in *traip*^-^ CB-NBs via *wor-GAL4*, including Full Length (FL), RING mutant (I8D, A10E), and several truncations and internal deletions based on the major features of Traip (Figure S5A). GFP::Traip^FL^ recapitulated the interphase and mitotic localizations of the mNG::Traip CRISPR transgenic (Figure 7D and S5B). GFP::Traip^I8D,A10E^ localized to the proper sites, but formed some aggregates (Figure S5C). Neither the RING domain nor the first coiled coil were sufficient to mediate any localization (Figure S5D). Both the second coiled coil and the C-terminal region were sufficient to mediate centrosome localization (Figure S5E-S5H). Finally, the C-terminal region was both necessary and sufficient for localization to the furrow during mitosis and the nucleus/nucleolus during interphase (Figure S5G and S5H). We also tested whether these protein regions are required for Traip MB function, but none were sufficient to rescue *traip*^-^ MB size (Figure S6A) indicating that Traip function requires multiple domains. Human GFP::TRAIP also failed to rescue *traip^-^* MB size (Figure S6A).

Reasoning that Traip could control MB development through either an interphase or a mitotic function, we first tested whether Traip has any major DDR functions in interphase. Using γH2Av staining to identify double stranded DNA breaks in 24 hours APF MB-NBs, we found that control (Figure S7A) and most *traip*^-^ (Figure S7B) MB-NBs had few γH2Av puncta. A small subset of *traip*^-^ MB-NBs had extremely elevated γH2Av puncta (Figure S7C); however, these cells appeared to be apoptotic, and thus likely reflected an indirect effect of *traip^-^* (caspase-dependent cell death) rather than a direct effect of impaired interphase DNA repair. Excluding these outliers, γH2Av puncta numbers were not different between controls and *traip*^-^ (Figure S7D). We also hypothesized that if Traip functions in interphase DNA repair, then activation of DNA damage signaling might potentiate cell death in *traip*^-^ MB-NBs. However, knocking down *mei-41*, *chk1*, or *p53* via RNAi did not significantly improve *traip*^-^ MB size (Figure S7E). These data suggest that interphase DNA repair is not a major function for *Traip* in MB-NBs under normal conditions.

We next tested whether Traip primarily controls MB development through a mitotic function. GFP::Traip^ΔNLS^, which contains a deletion of the nuclear localization signal (NLS), localized similarly to wild type GFP::Traip during mitosis but was cytoplasmic in interphase (Figure 7F). Nonetheless, *GFP::Traip^ΔNLS^* fully rescued *traip*^-^ MB size (Figure 7G and 7H), indicating that a mitotic Traip function is critical for MB-NBs. To test whether proper mitotic localization is important for Traip function, we introduced a Tom20 tag to ectopically force Traip to the mitochondria. Tom20::TagRFP::Traip localized cytoplasmically in interphase, and was absent from the spindle region in mitosis (Figure 7I). Mitochondria-localized Tom20::TagRFP::Traip provided an intermediate rescue of *traip*^-^ MB size (Figure 7H and 7J). Thus, proper mitotic localization of Traip to the spindle, centrosome, and/or midbody is important for full Traip function.

## Discussion

Our study in *Drosophila* shows that *traip*^-^ shares several characteristics with human microcephaly mutants. First, the *traip*^-^ phenotype is highly brain-specific, with body defects being extremely rare. Second, the *traip*^-^ MB phenotype is developmental rather than neurodegenerative in nature, reflecting a primary rather than secondary microcephaly-like disorder. Finally, as with many human microcephaly genes, *Traip* functions to promote neural progenitor cell (NPC) proliferation and survival. Thus, *traip*^-^ represents a powerful new disease model for understanding the etiological mechanisms underlying microcephaly.

Despite their ubiquitous expression, mutations in microcephaly genes primarily affect the cerebral cortex in humans. Similarly, both *Traip* and the DDR microcephaly gene *MCPH1* (Rickmyre et al., 2007) are ubiquitously expressed in *Drosophila*, yet the MB is the only adult structure affected in their mutants. While most NBs have a limited window of proliferation, MB-NBs divide continuously from embryogenesis into late pupal stages (Ito and Hotta, 1992; Truman and Bate, 1988), potentially allowing more accumulation of rare or small effects over many cell cycles. Interestingly, our drug experiments indicate that *traip*^-^ mitotic DNA bridges may be the result of inter-strand crosslinks rather than late replicating regions; perhaps the highly proliferative MB-NBs generate increased reactive oxygen species and aldehydes, leading to increased inter-strand crosslinks. Additionally, while many tissues can make up for lost cells via compensatory proliferation (Haynie and Bryant, 1977; Pfau et al., 2016), no such process appears to exist for replacing lost NPCs. Thus, we speculate that the MBs are especially sensitive to mutations in microcephaly genes as a consequence of rapid, prolonged proliferation and inability to compensate for lost progenitor cells, and that a similar explanation may account for the sensitivity to microcephaly gene mutation in the human cortex.

Our work provides the first link between a known function of Traip and proper brain development. We found that interphase nuclear localization is not required for Traip function, interphase double stranded DNA breaks are not increased in *traip*^-^ MB-NBs, and DNA damage signaling does not appear to mediate *traip*^-^ MB-NB death, and thus conclude that Traip interphase functions are dispensable for MB-NB survival under normal conditions. Instead, we discovered the presence of mitotic DNA bridges, sensitivity to inter-strand crosslinking agents, RING domain-dependence, and requirement for mitotic localization only, and therefore conclude that the primary function for Traip in MB-NBs is to ubiquitylate and remove stalled replication machinery during mitosis (Figure 8A; Deng et al., 2019; Priego Moreno et al., 2019; Sonneville et al., 2019). Thus, we surmise that *traip*^-^MB-NBs have stalled replication machinery which remains loaded throughout mitosis, preventing mitotic DNA synthesis repair and proper sister chromatid segregation in mitosis (Figure 8B). As anaphase proceeds, attached sister chromatids are pulled to opposite poles and they form UFBs as the under-replicated DNA is stretched out between them. These bridges could be physically broken, leading to chromosome fragmentation, generating aneuploidy or micronuclei and causing nuclear deformations in daughter cells (Gisselsson et al., 2001; Heddle and Carrano, 1977). Alternatively, persistence of DNA bridges at the cytokinetic furrow could induce mitotic exit and furrow regression (Pampalona et al., 2012; Shi and King, 2005), leading to multiple nuclei or polyploidy which likely prevent further proliferation.

**Figure 8.**
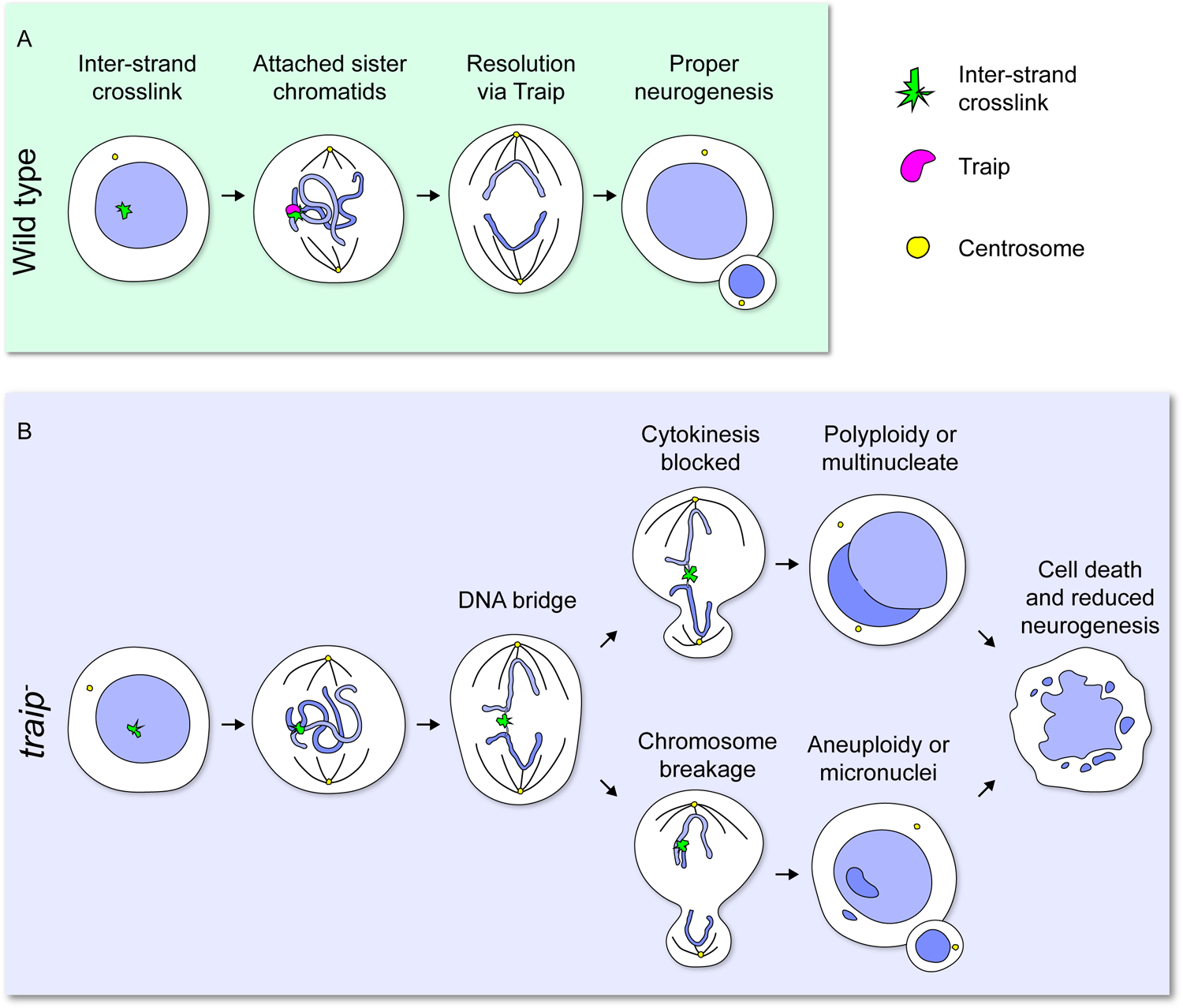
Model of Traip function in neurogenesis. (A) In wild type, inter-strand crosslinks may prevent completion of DNA replication, resulting in sister chromatids remaining partially attached in mitosis. Traip initiates the resolution of these attached sister chromatids, thereby ensuring chromosome segregation and promoting proper neurogenesis. (B) In *traip^-^*, attached sister chromatids are not properly resolved, leading to DNA bridges in mitosis. DNA bridges could block cytokinesis, leading to formation of polyploid or multinucleated cells (top path). Alternatively, DNA bridges could break during mitosis, leading to formation of aneuploid cells or micronuclei (bottom path). These defects could result in cell cycle exit, apoptosis, and ultimately reduced neurogenesis.

We found that *traip*^-^MB-NBs are prematurely lost via caspase-dependent cell death and thus fail to generate proper KC numbers. DNA bridge-induced defects likely feed into apoptosis, but further work is required to dissect the pathways connecting them. In *Drosophila*, polyploid NBs can accumulate significant DNA damage as they enter mitosis (Nano et al., 2019), and chromosome breakage during mitosis in *traip*^-^could induce death through DNA damage signaling. *Drosophila* embryos laid by *traip*^-^mothers are lethal, with extensive chromosome bridging and *Chk2*-dependent cell death, suggesting DNA damage accumulation leads to apoptosis in the rapidly dividing cells of the early embryo (Merkle et al., 2009). However, our RNAi experiments in the MB indicate that DNA damage signaling is not a major contributor to MB-NB apoptosis (Figure S7E). In mammalian NPCs, polypoidy and binucleation can cause G1 arrest and apoptosis (Aylon and Oren, 2011; Storchova and Kuffer, 2008). In *Drosophila*, neurons can become polyploid in response to DNA damage (Nandakumar et al., 2020), and NBs can become massively polyploid in some mutants (Poulton et al., 2017; Swider et al., 2019), suggesting that, even though polyploidy may be better tolerated in flies, polyploid NBs are unlikely to successfully complete additional mitoses. We infer the existence of *traip*^-^aneuploid MB-NBs which produce a wide range of daughter KC numbers, suggesting that *traip*^-^generates some aneuploidies that are well-tolerated and others that are highly lethal to the cell. This parallels the situation in mammals, where aneuploid NPCs and neurons are common (Rehen et al., 2001) but also sensitive to G1 arrest, cell cycle exit, and apoptosis (Peterson et al., 2012; Storchova and Kuffer, 2008). Thus, both polyploidy and aneuploidy could stop further proliferation in *traip*^-^MB-NBs by preventing proper mitosis or inducing G1 arrest and cell cycle exit, eventually triggering apoptosis.

In our study we identified a centrosome, spindle, and cytokinetic furrow localization for Traip which is important for function. One possibility is that the dynamic movement of Traip on the mitotic spindle and cytokinetic furrow promotes encounters with unresolved DNA bridges. We never observed GFP::Traip on bridges; however, it is thought that a single TRAIP protein is sufficient to unload each replisome (Wu et al., 2019), so fluorescence detection may be unlikely. Interestingly, centrosome localization is a common aspect of microcephaly-linked proteins, including MCPH1 which also functions in DDR (Jeffers et al., 2008; Rai et al., 2008). Similar to Traip, MCPH1 has mitotic functions required for proper chromosome segregation, and mutations in *MCPH1* lead to lagging chromosomes, DNA bridges, and micronuclei (Arroyo et al., 2017; Rickmyre et al., 2007). Mutations in microcephaly genes with centrosome-associated functions, such as *CEP135* and CDK5RAP2, cause dysregulation of centrosome numbers which also lead to chromosome segregation errors and aneuploidy (Barrera et al., 2010; Hussain et al., 2012; Shi et al., 2019). Thus, mitotic roles, ensuring proper chromosome segregation, and suppressing aneuploidy are common features of microcephaly-linked proteins. Future work seeking to better understand these shared defects may reveal a deeper etiological connection across microcephaly disorders.

## Materials and Methods

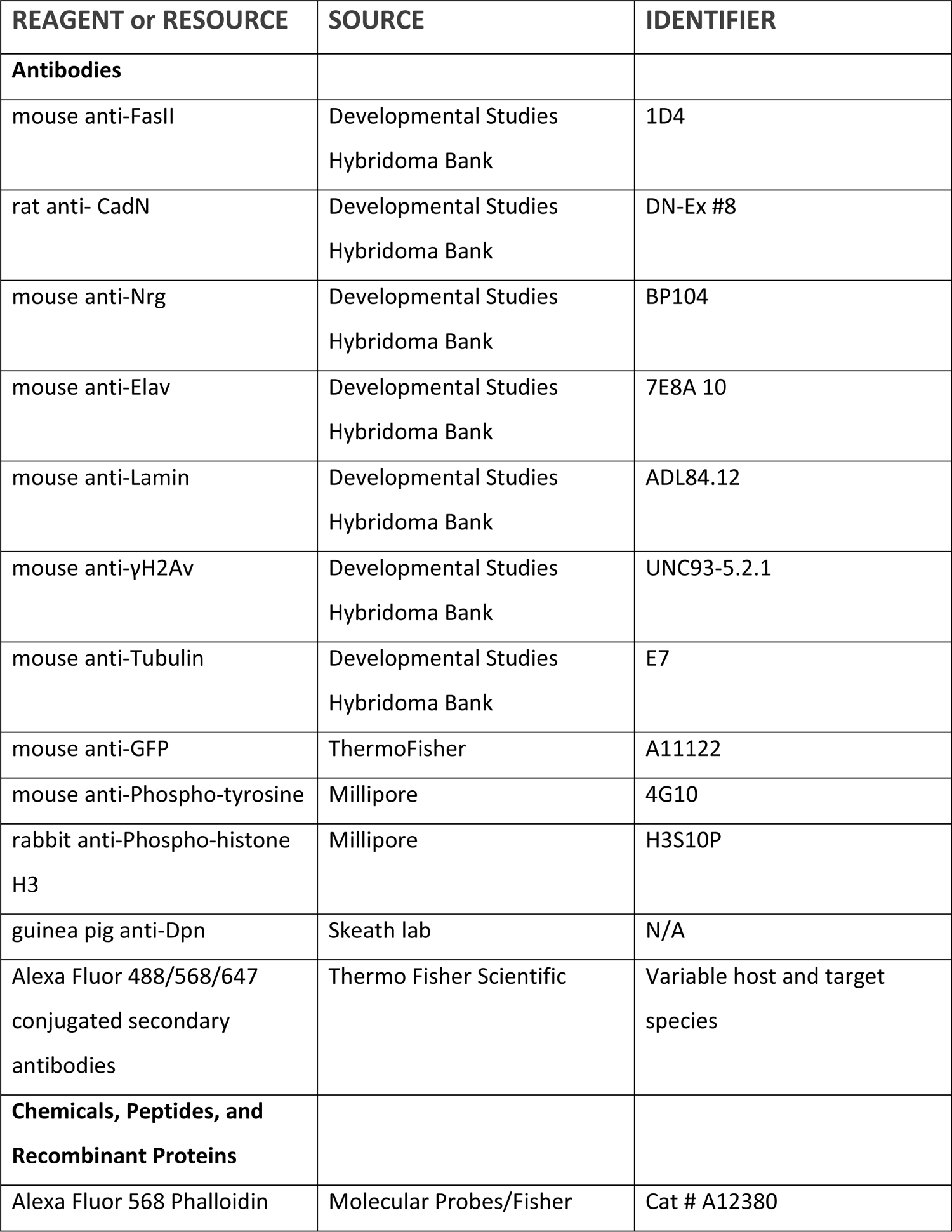
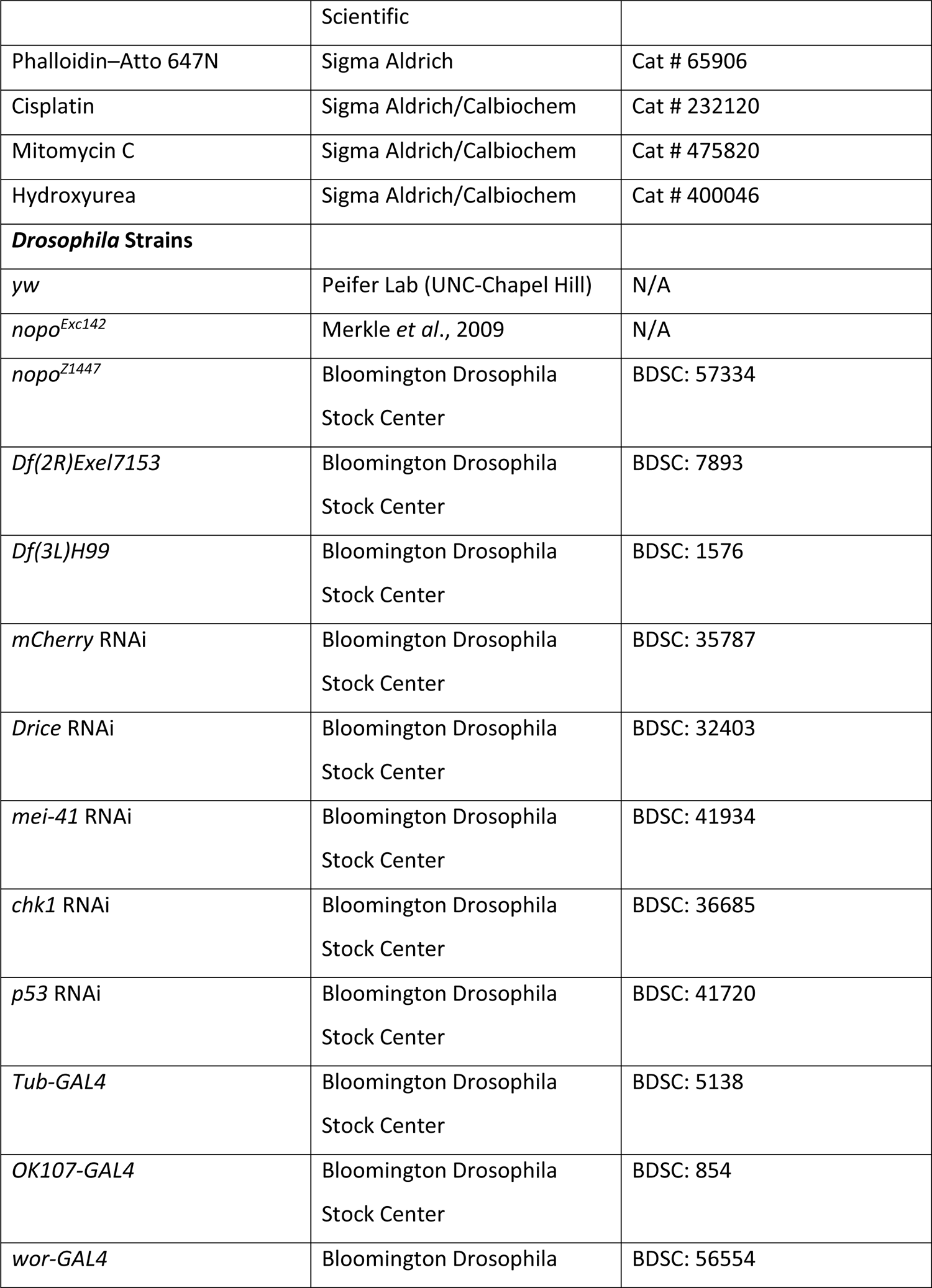
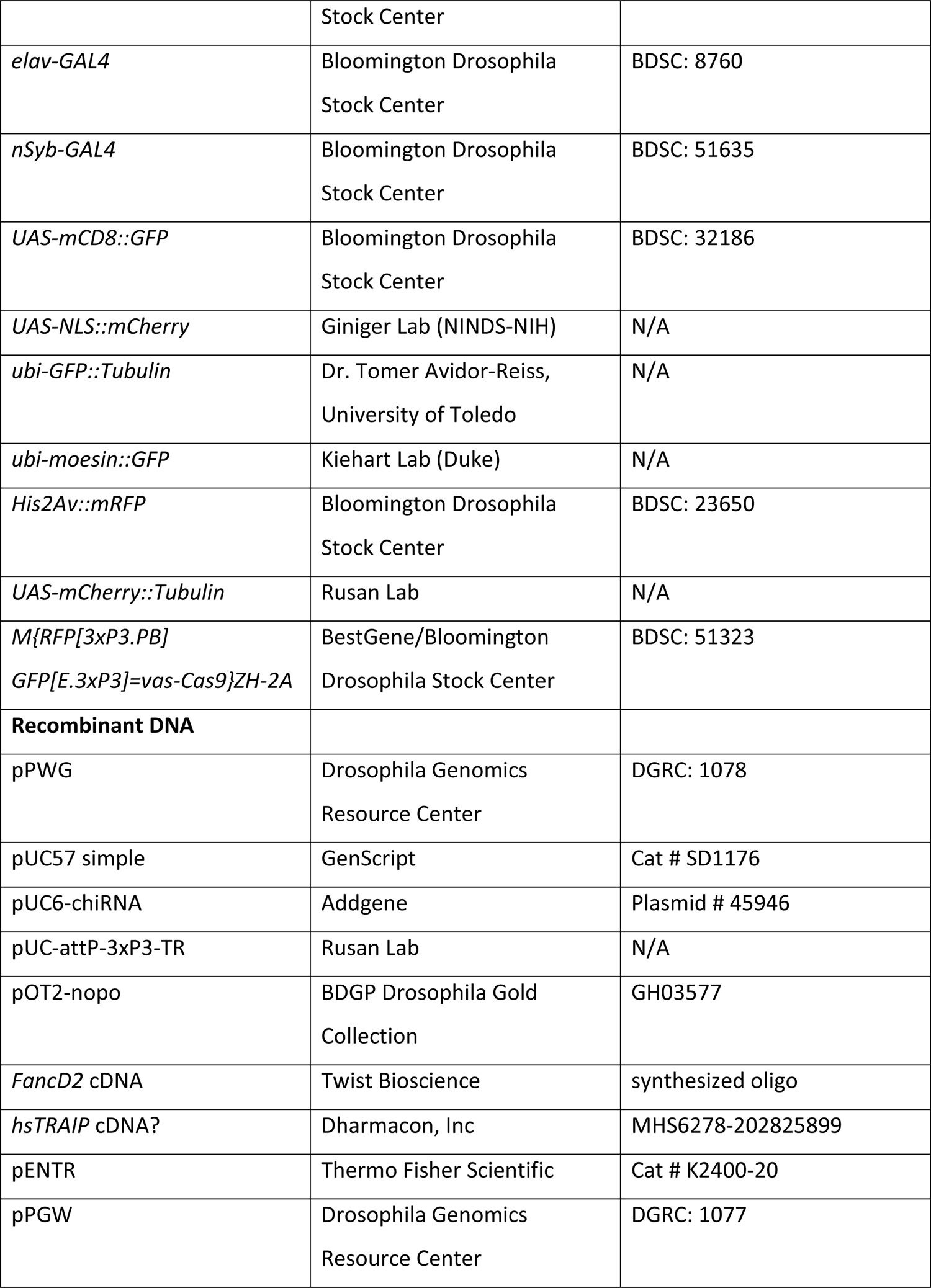
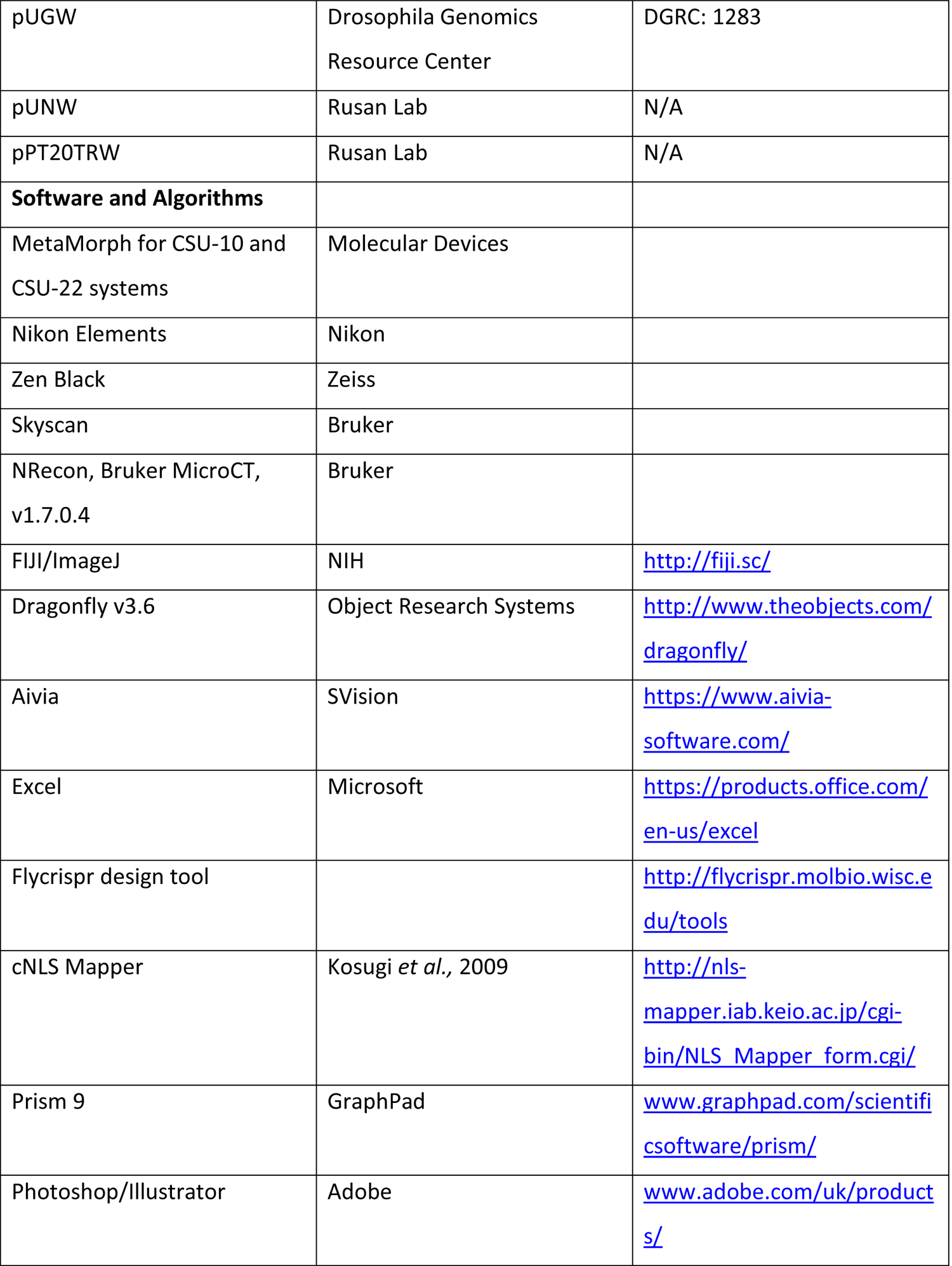

| REAGENT or RESOURCE | SOURCE | IDENTIFIER |
| --- | --- | --- |
| **Antibodies** |  |  |
| mouse anti-FasII | Developmental Studies<br>Hybridoma Bank | 1D4 |
| rat anti- CadN | Developmental Studies<br>Hybridoma Bank | DN-Ex #8 |
| mouse anti-Nrg | Developmental Studies<br>Hybridoma Bank | BP104 |
| mouse anti-Elav | Developmental Studies<br>Hybridoma Bank | 7E8A 10 |
| mouse anti-Lamin | Developmental Studies<br>Hybridoma Bank | ADL84.12 |
| mouse anti-γH2Av | Developmental Studies<br>Hybridoma Bank | UNC93-5.2.1 |
| mouse anti-Tubulin | Developmental Studies<br>Hybridoma Bank | E7 |
| mouse anti-GFP | ThermoFisher | A11122 |
| mouse anti-Phospho-tyrosine | Millipore | 4G10 |
| rabbit anti-Phospho-histone<br>H3 | Millipore | H3S10P |
| guinea pig anti-Dpn | Skeath lab | N/A |
| Alexa Fluor 488/568/647<br>conjugated secondary<br>antibodies | Thermo Fisher Scientific | Variable host and target<br>species |
| **Chemicals, Peptides, and**<br>**Recombinant Proteins** |  |  |
| Alexa Fluor 568 Phalloidin | Molecular Probes/Fisher | Cat # A12380 |
|  | Scientific |  |
| Phalloidin–Atto 647N | Sigma Aldrich | Cat # 65906 |
| Cisplatin | Sigma Aldrich/Calbiochem | Cat # 232120 |
| Mitomycin C | Sigma Aldrich/Calbiochem | Cat # 475820 |
| Hydroxyurea | Sigma Aldrich/Calbiochem | Cat # 400046 |
| ***Drosophila* Strains** |  |  |
| *yw* | Peifer Lab (UNC-Chapel Hill) | N/A |
| *nopo*<sup>Exc142</sup> | Merkle *et al.*, 2009 | N/A |
| *nopo*<sup>Z1447</sup> | Bloomington Drosophila<br>Stock Center | BDSC: 57334 |
| *Df(2R)Exel7153* | Bloomington Drosophila<br>Stock Center | BDSC: 7893 |
| *Df(3L)H99* | Bloomington Drosophila<br>Stock Center | BDSC: 1576 |
| *mCherry* RNAi | Bloomington Drosophila<br>Stock Center | BDSC: 35787 |
| *Drice* RNAi | Bloomington Drosophila<br>Stock Center | BDSC: 32403 |
| *mei-41* RNAi | Bloomington Drosophila<br>Stock Center | BDSC: 41934 |
| *chk1* RNAi | Bloomington Drosophila<br>Stock Center | BDSC: 36685 |
| *p53* RNAi | Bloomington Drosophila<br>Stock Center | BDSC: 41720 |
| *Tub-GAL4* | Bloomington Drosophila<br>Stock Center | BDSC: 5138 |
| *OK107-GAL4* | Bloomington Drosophila<br>Stock Center | BDSC: 854 |
| *wor-GAL4* | Bloomington Drosophila | BDSC: 56554 |
|  | Stock Center |  |
| *elav-GAL4* | Bloomington Drosophila<br>Stock Center | BDSC: 8760 |
| *nSyb-GAL4* | Bloomington Drosophila<br>Stock Center | BDSC: 51635 |
| *UAS-mCD8::GFP* | Bloomington Drosophila<br>Stock Center | BDSC: 32186 |
| *UAS-NLS::mCherry* | Giniger Lab (NINDS-NIH) | N/A |
| *ubi-GFP::Tubulin* | Dr. Tomer Avidor-Reiss,<br>University of Toledo | N/A |
| *ubi-moesin::GFP* | Kiehart Lab (Duke) | N/A |
| *His2Av::mRFP* | Bloomington Drosophila<br>Stock Center | BDSC: 23650 |
| *UAS-mCherry::Tubulin* | Rusan Lab | N/A |
| *M{RFP[3xP3.PB]*<br>*GFP[E.3xP3]=vas-Cas9}ZH-2A* | BestGene/Bloomington<br>Drosophila Stock Center | BDSC: 51323 |
| **Recombinant DNA** |  |  |
| pPWG | Drosophila Genomics<br>Resource Center | DGRC: 1078 |
| pUC57 simple | GenScript | Cat # SD1176 |
| pUC6-chiRNA | Addgene | Plasmid # 45946 |
| pUC-attP-3xP3-TR | Rusan Lab | N/A |
| pOT2-nopo | BDGP Drosophila Gold<br>Collection | GH03577 |
| *FancD2* cDNA | Twist Bioscience | synthesized oligo |
| *hsTRAIP* cDNA? | Dharmacon, Inc | MHS6278-202825899 |
| pENTR | Thermo Fisher Scientific | Cat # K2400-20 |
| pPGW | Drosophila Genomics<br>Resource Center | DGRC: 1077 |
| pUGW | Drosophila Genomics Resource Center | DGRC: 1283 |
| pUNW | Rusan Lab | N/A |
| pPT20TRW | Rusan Lab | N/A |
| **Software and Algorithms** |  |  |
| MetaMorph for CSU-10 and CSU-22 systems | Molecular Devices |  |
| Nikon Elements | Nikon |  |
| Zen Black | Zeiss |  |
| Skyscan | Bruker |  |
| NRecon, Bruker MicroCT, v1.7.0.4 | Bruker |  |
| FIJI/ImageJ | NIH | http://fiji.sc/ |
| Dragonfly v3.6 | Object Research Systems | http://www.theobjects.com/dragonfly/ |
| Aivia | SVision | https://www.aivia-software.com/ |
| Excel | Microsoft | https://products.office.com/en-us/excel |
| Flycrispr design tool |  | http://flycrispr.molbio.wisc.edu/tools |
| cNLS Mapper | Kosugi *et al.*, 2009 | http://nls-mapper.iab.keio.ac.jp/cgi-bin/NLS_Mapper_form.cgi/ |
| Prism 9 | GraphPad | www.graphpad.com/scientificsoftware/prism/ |
| Photoshop/Illustrator | Adobe | www.adobe.com/uk/products/ |

### D. melanogaster

Experimental fly crosses were maintained on Bloomington Recipe Fly food from LabExpress (Ann Arbor, MI) and kept at 25°C. Crosses were either 8 virgin females per vial or 20 virgin females per bottle, with at least half as many males. *yw* was used as a control in all genotypes marked “+”. All new transgenic animals were generated using standard embryo injection protocols by BestGene (Chino Hills, CA).

### Generation of Transgenic Drosophila

For *traip^Δ^*, homology arms were cloned into pUC-attP-3xP3-TR, such that the repair construct contained 1100 bp upstream of the *Traip* start codon and 1094 bp downstream of the *Traip* stop codon flanking a attP-3xP3-TR replacement. For *mNG::Traip*, homology arms and mNeonGreen were cloned into pUC57, such that the repair construct contained 2226 bp upstream of the *Traip* start codon, the mNeonGreen coding sequence, and 1093 bp downstream of the *Traip* start codon. Guide RNAs were cloned into pU6-chiRNA. Note that PAM sequences in homology arms were mutated such that they were not predicted to be targets Cas9. Repair template and gRNA plasmid constructs were injected into *y^1^ M{RFP[3xP3.PB] GFP[E.3xP3]=vas-Cas9}ZH-2A w^1118^/FM7c* using standard procedures. Putative CRISPR-positive alleles were crossed to *CyO-Cre* to eliminate 3xP3-TagRFP and were fully sequenced from outside the homology arms. *traip^Δ^* was further back-crossed to *yw* for three generations and re-sequenced to eliminate a second site lethal mutation. Both CRISPR alleles were crossed to *Df(2R)Exel7153* to test for maternal effect lethality; *traip^Δ^* was maternal effect lethal, and *mNG::Traip* was fertile.

*Traip^I8D,A10E^* and *Traip^L12E,G14D^* were rationally designed using amino acid alignment of Traip and TRAF6, which is a RING domain E3 ligase that has a known crystal structure with its E2 conjugating enzyme Ubc13 (Yin et al 2009). *Traip^ΔNLS^* was designed using cNLS Mapper, which predicts importin α-dependent, CDK1-regulated NLS sequences (Kosugi et al 2009). *Traip^I8D,A10E^*, *Traip^L12E,G14D^* and *Traip^ΔNLS^* (deletion of positions 357-366 YSIFKKPRLL) were generated using mutagenic primers. *Traip* truncations were PCR amplified from cDNA. *Traip*, *Traip* variants, human *TRAIP*, and *FancD2* cDNAs were cloned into pENTR using standard Gateway cloning methods. Gateway cloning was used to move constructs from pENTR into pPGW for *UAS-GFP::Traip^variant^*^s^, pUGW for *ubi-GFP::Traip*, pPT20TRW for *UAS-Tom20::TagRFP::Traip*, and pUNW for *ubi-mNG::FancD2*.

### Drug Treatments

Parent flies (*traip^Exc142^*/*CyO* females and either *yw* or *traip^Δ^* males) were crossed for 2-3 days and then transferred to fly food made from 0.6 g of Carolina Formula 4-24 (Fisher Scientific) and 2 mL of ddH_2_O. Parents laid eggs for 24 hours, eggs were allowed to hatch for 24 hours, and then 450 μL of yeast solution (1 g dry yeast sprinkles per 10 mL ddH_2_O) with drug was added on top of the food. Drugged larvae were allowed to develop and adults were scored by genotype daily as they eclosed. Concentrated drug stock solutions were reconstituted as follows: Cisplatin at 1 mg/mL in phosphate buffered saline (PBS); Mitomycin C at 0.5 mg/mL in ddH_2_O; Hydroxyurea at 5 mg/mL in ddH_2_O.

### μ-CT

Control, *traip*^-^, and *traip*^-^ + *ubi-GFP::Traip* flies were processed for μ-CT according to Schoborg *et al.,* 2019. Briefly, adult flies were aged to 2-4 days old, anesthetized on CO_2_, de-waxed in PBST (0.5% Triton X-100), and fixed in Bouin’s solution for 24 hours. Fixed flies were washed in 0.1 M Na_2_HPO_4_ with 1.8% sucrose and then stained in 0.1 N solution of I_2_KI for 2 days. Stained flies were washed with ultrapure H_2_O and scanned using SkyScan 1172 desktop scanner. Tomograms were generated with NRecon, and 3D volumetric segmentation was performed using Dragonfly. Brain measurements were normalized to thorax width to control for overall size differences among individual flies.

### Immunostaining

Animals at appropriate stages were collected and aged: adult flies were aged to 3-4 days old unless otherwise stated; white pre-pupae were collected and aged for 24, 48, or 72 hours to obtain pupal stages. Staining procedures were similar to Janelia FlyLight protocols (https://www.janelia.org/project-team/flylight/protocols). Tissues were dissected in SF900 S2 cell media (Fisher Scientific) and fixed with 2% paraformaldehyde in SF900 for 2 hours at room temperature (RT) or overnight at 4°C. Fixed samples were washed three times in PBST (PBS + 0.5% Triton-X100) and blocked for 1.5 hours in PBST + 5% normal goat serum (NGS). Blocked samples were incubated in PBST + NGS with primary antibody for 1-2 hours at RT and then 1-2 overnights at 4°C. Primary antibody concentrations were as follows: mouse anti-FasII (DSHB 1D4) 1:50; rat anti-CadN (DSHB DN-Ex#8) 1:25; mouse anti-Nrg (DSHB BP104) 1:25; mouse anti-Elav (DSHB 7E8A 10) 1:100; mouse anti-Lamin (DSHB ADL84.12) 1:100; mouse anti-γH2Av (DSHB UNC93-5.2.1) 1:100; mouse anti-Tubulin (DSHB E7) 1:200; mouse anti-GFP (Thermo Fisher A11122) 1:1000; mouse anti-Phospho-tyrosine (Millipore 4G10) 1:100; rabbit anti-Phospho-histone H3 (Millipore H3S10P) 1:40000; guinea pig anti-Dpn (Skeath lab, Washington University in St. Louis) 1:50. Samples were washed five times in PBST + NGS, and then incubated in PBST + NGS with 1:500 secondary antibody (Thermo Fisher) and 1x DAPI for 1-2 hours at RT and then 1-3 overnights at 4°C. Samples were washed three times in PBST + NGS and then three times in PBST, before being post-fixed in 4% paraformaldehyde in PBS for 1 hour at RT. Post-fixed samples were washed twice in PBST, and then washed in PBS until all Triton-X100 was removed. Samples were mounted on poly-L-lysine coated #1.5 coverslips, rinsed in ddH_2_O, and then dehydrated through a series of 30%, 50%, 75%, 95% and then three 100% ethanol 10 minute baths. Dehydrated coverslips were then bathed three times in 100% xylene for 5 minutes, before being mounted in DPX on slides with #1.5 coverslips as spacers. DPX was cured at RT overnight. For phalloidin-stained samples, Phalloidin–Atto 647N (Sigma Aldrich) was added at 1:40 in PBS for 20 minutes after secondary antibody incubation, and samples were not post-fixed and were mounted in AquaPoly Mount (Polysciences, Inc.).

### Light Microscopy

Most fixed samples were imaged using a Zeiss LSM 880 Confocal Microscope with a 63x/1.4 NA objective, GaAsP detectors, and 405, 488, 561, and 641 nm laser lines, controlled using Zen Black software. Some fixed samples (Figures 4A, 5A, 5I, 5J) were imaged using a Nikon W1 spinning disc confocal equipped with a Prime BSI cMOS camera (Photometrics) and a 100X/1.4 NA silicon immersion objective, controlled using Nikon Elements software. Live imaging was performed on either using the same Nikon W1 with 100x/1.4 NA silicon or 40x/1.3 NA oil immersion objective, or using an Eclipse Ti2 (Nikon) with a 100X/1.49 NA objective or a 40x/1.3 NA objective and a 1.5X tube lens, an ORCA-Flash 4.0 CMOS camera (Hamamatsu Photonics), and 405, 491, 561, and 642 nm laser lines, controlled by MetaMorph software. For live imaging, larval brains were dissected in Schneider’s media and mounted by sandwiching between a #1.5 coverslip and a 50-mm lummox dish, with droplets of Halocarbon oil 700 as a cushion and surrounding the edge of the coverslip. ImageJ was used to generate maximum intensity projections for presentation.

### Image Analysis

For MB α lobe cross-sectional analysis, image stacks of α lobes were stack rotated to be perpendicular to the viewing plane using the Interactive Stack Rotation plugin in ImageJ. Rotated MB lobes were then cross sectioned through the middle of the lobe using the reslice tool, and the area of the resliced MB lobe was measured in ImageJ. For MB volumetric analysis, either FasII-positive or *OK107-GAL4* > mCD8::GFP-positive MB lobes were segmented and measured using the Pixel Classification and 3D Object Analysis tools in Aivia software. For KC counting analysis, the *OK107-GAL4* > NLS::mCherry-positive KCs were similarly segmented and counted using Pixel Classification and 3D Object Analysis tools in Aivia software. γH2Av puncta were thresholded and identified using the 3D Objects Counter in ImageJ. Then, γH2Av objects inside the nucleus, as marked by anti-lamin immunostaining, were counted in each MB-NB.

### Statistical analysis

Data analysis was performed using Microsoft Excel and GraphPad Prism. In all graphs, the mean ± standard deviation, and all individual data points are presented. Sample sizes were primarily determined based on the availability of animals of the proper genotype and developmental stage, and the time required for dissection, processing, and imaging; given these considerations, we processed 8-12 brains per genotype for most experiments, although some brains were not imaged due to physical damage. Experiments measuring MB size were typically performed once, with control and *traip^Δ^* conditions repeated in each experiment. Shapiro-Wilk test was used to test the assumption of normality. Statistical tests used to determine significance include t-test, Mann-Whitney U test, ANOVA with Tukey’s test, and chi-squared test, and are reported in Figure legends.

## Acknowledgements

We thank the National Heart, Lung, and Blood Institute Light Microscopy Core, especially Xufeng Wu, for support with confocal microscopy. We thank Bloomington Drosophila Stock Center for fly stocks, and Developmental Studies Hybridoma Bank for antibodies. Carey Fagerstrom performed all cloning. Rachel Ng performed the μ-CT experiments. We thank Alex Kelly, Matthew Hannaford, and Ramya Varadarajan for helpful discussion, and Todd Schoborg, Brian Galletta, Ed Giniger, Hong Xu for critically reading the manuscript. This work is supported by the Division of Intramural Research at the NHLBI/NIH (1ZIAHL006126 to N.M.R.).

## Author Contributions

R.S.O. conceived of the project, designed and performed all experiments, and wrote the manuscript. N.M.R. oversaw the project, designed and discussed experiments, and edited the manuscript.

---

**Figure S1.**
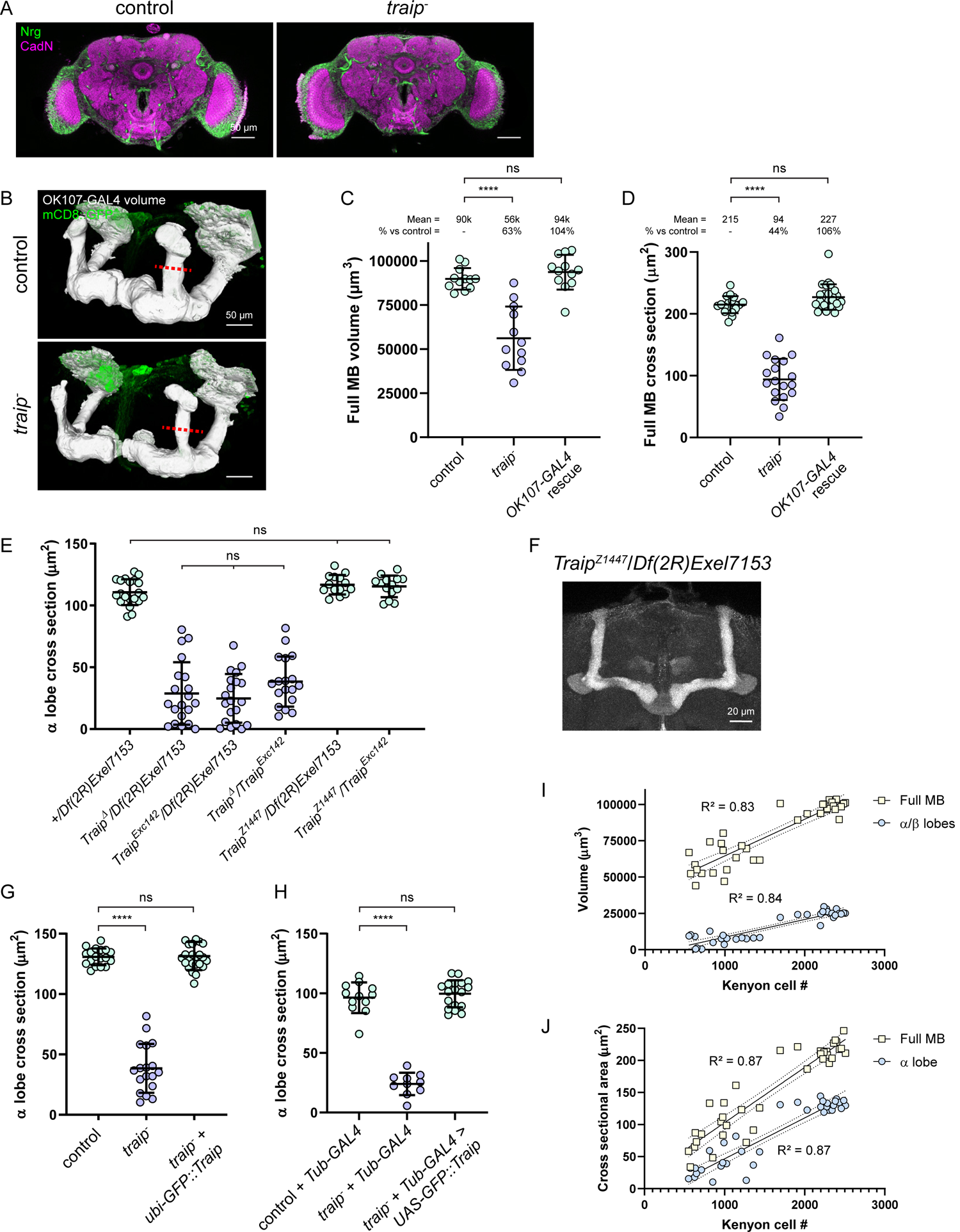
*Traip* is required for proper MB structure, supporting figures. (A) Confocal slices through control and *traip^-^* adult brains stained for N-Cadherin (CadN, magenta) and Neuroglian (Nrg, green), showing no obvious differences in neuropil regions or axon tracts, excluding the MBs. (B) Control and *traip^-^* full MB volumes (white) segmented from *OK107-GAL4* > mCD8::GFP fluorescence (green). (C) Full MB volume measurements. N ≥ 18 MBs. (D) αˈ/α lobe cross-section measurements. N ≥ 18 MBs. (E) α lobe cross-section measurements of controls and various combinations of *Traip* mutant alleles. Homozygous null animal MBs are similarly reduced, whereas hypomorphic animals have wild-type MB size. N ≥ 18 MBs/genotype. (F) Hypomorphic *traip^Z1447^* MBs have wild-type morphology. (G, H) α lobe cross-section measurements show that *traip^-^* + *ubi-GFP::Traip* (G) and *traip^-^* + *Tub-GAL4 > GFP::Traip* (H) rescues have wild-type MB size. N ≥ 10 MBs. (I). Simple linear regression between KC number (X axis) and volumes (Y axis) of *OK107-GAL4* > mCD8::GFP-positive full MBs (R^2^ = 0.83, yellow squares) or FasII-positive α/β lobes (R^2^ = 0.84, blue circles). (J) Simple linear regression between KC number (X axis) and the cross sectional areas (Y axis) of *OK107-GAL4* > mCD8::GFP-positive αˈ/α lobes (R^2^ = 0.87, yellow squares) or FasII-positive α lobes (R^2^ = 0.87, blue circles). Two-tailed Mann-Whitney test was used for significance. ns = not significant, **** p < 0.0001. Scale bars = 50 μm (A, B), 20 μm (F).

**Figure S2.**
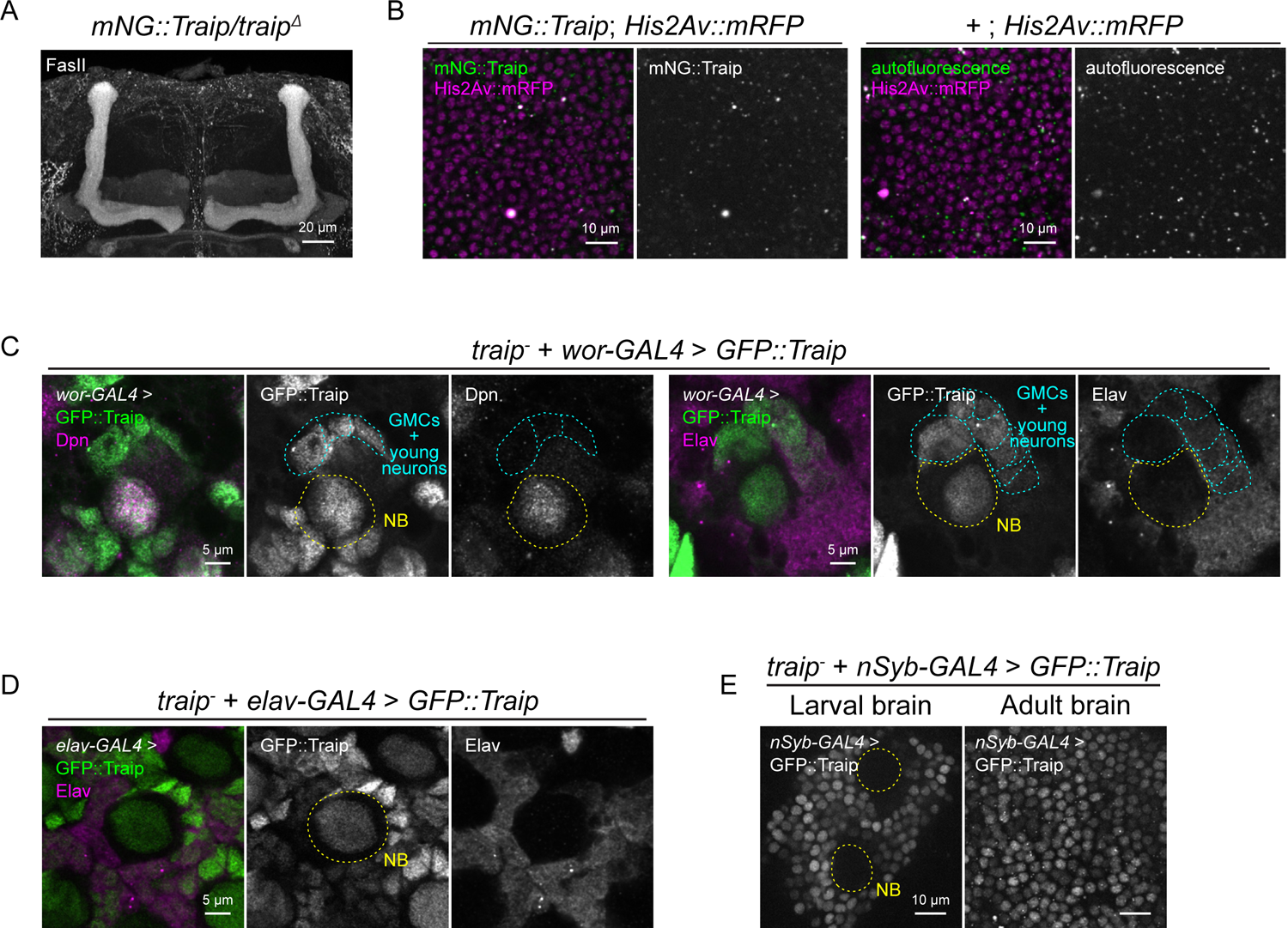
*Traip* is required in neuroblasts, supporting figures. (A) *mNG::Traip^CRISPR^* encodes a functional protein that fully rescues *traip^Δ^* MB size. (B) Adult brains from either *mNG::Traip^CRISPR^* (left panels) or *+* (untagged *Traip*, right panels) with His2Av::mRFP (magenta). mNG::Traip (green, gray; left panels) does not have fluorescent signal above autofluorescence background of *+* (green, gray; right panels). (C) CB-NBs from *traip^-^* + *wor-GAL4* > *GFP::Traip* 3^rd^ instar larval brains, stained either Dpn (magenta; left panels) or Elav (magenta; right panels). *wor-GAL4* > GFP::Traip (green) is expressed in NBs (Dpn-positive, yellow highlighting) and persists into daughter GMCs and neurons (Elav-positive, cyan highlighting). (D) *traip^-^* + *elav-GAL4* > *GFP::Traip* has significant GFP::Traip expression in 3^rd^ instar larval CB-NBs. (E) *traip^-^* + *nSyb-GAL4* > *GFP::Traip* has GFP::Traip expression in larval and adult neurons, but no expression in larval NBs. Scale bars = 10 μm (B, E), 5 μm (C, D).

**Figure S3.**
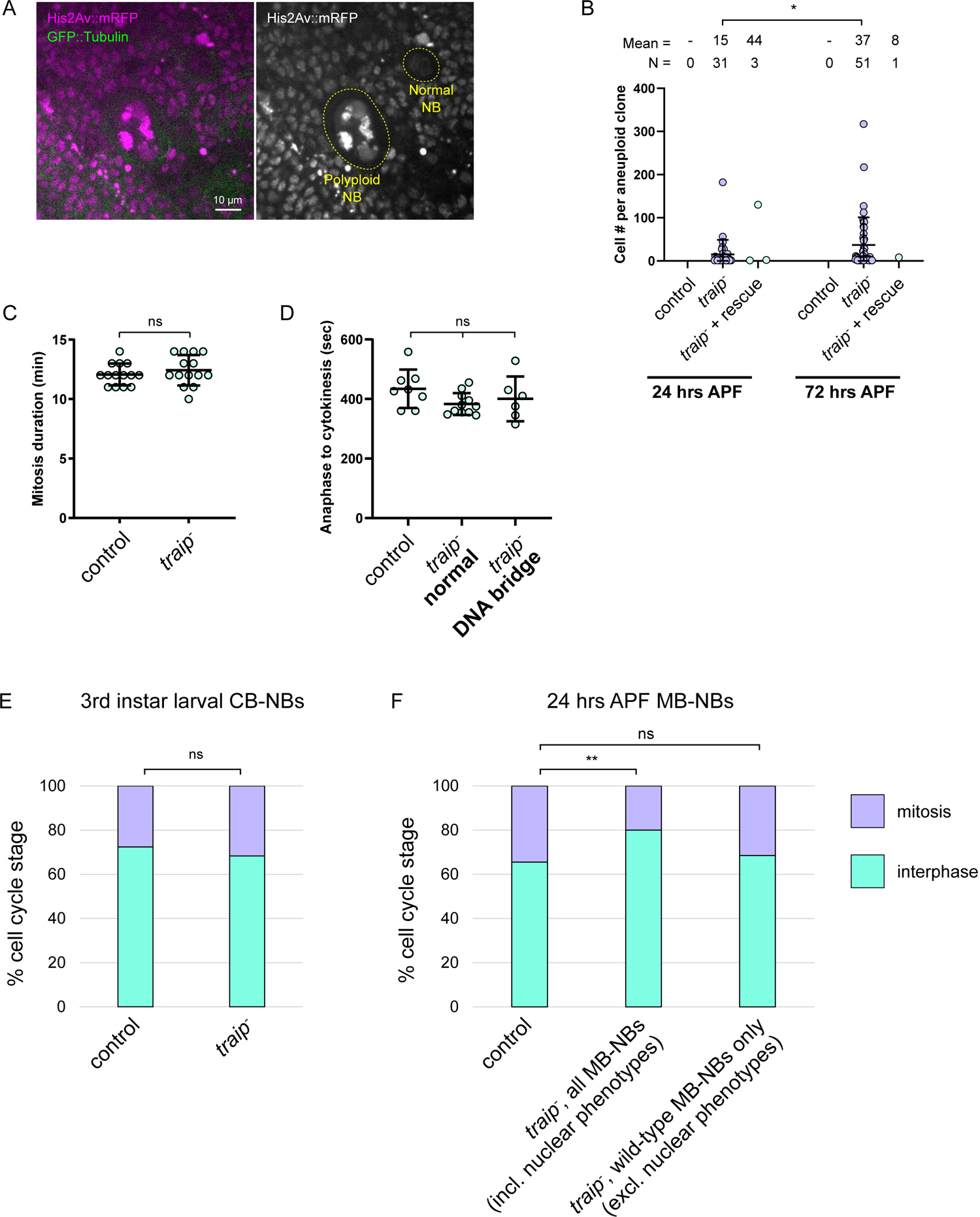
*Traip* suppresses multinuclear phenotypes and mitotic DNA bridges, supporting figures. (A) Live imaging of *traip^-^* 3^rd^ instar larval CB-NBs expressing His2Av::mRFP (magenta) and GFP::Tubulin (green) showing likely polyploid NBs with enlarged nuclei and increased His2Av::mRFP fluorescence compared to a normal NB. Scale bars = 10 μm. (B) KC number per aneuploid clone in control, *traip^-^*, and *traip^-^* + *GFP::Traip* rescue expressing *OK107-GAL4* > NLS::mCherry + mCD8::GFP at 24 and 72 hours APF pupal brains. The average number of KCs per clone in *traip^-^* increases from 15 at 24 hours APF to 37 at 72 hours APF. * p = 0.0287. (C) Total duration of mitosis for control and *traip^-^* 3^rd^ instar larval CB-NBs, as measured from prophase onset to full furrow constriction. N = 14 NBs. (D) Time from anaphase onset to full furrow constriction for control CB-NBs and *traip^-^* CB-NBs either with or without DNA bridges. (E) Mitotic index of fixed control and *traip^-^* 3^rd^ instar larval CB-NBs. (F) Mitotic index of fixed control and *traip^-^* 24 hours APF MB-NBs, either including all *traip^-^* MB-NBs or only *traip^-^* MB-NBs without nuclear phenotypes (right column). ** p = 0.0025. Two-tailed Mann-Whitney test (B), t-test (C, D), or chi-squared test (E, F) were used for significance. ns = not significant.

**Figure S4.**
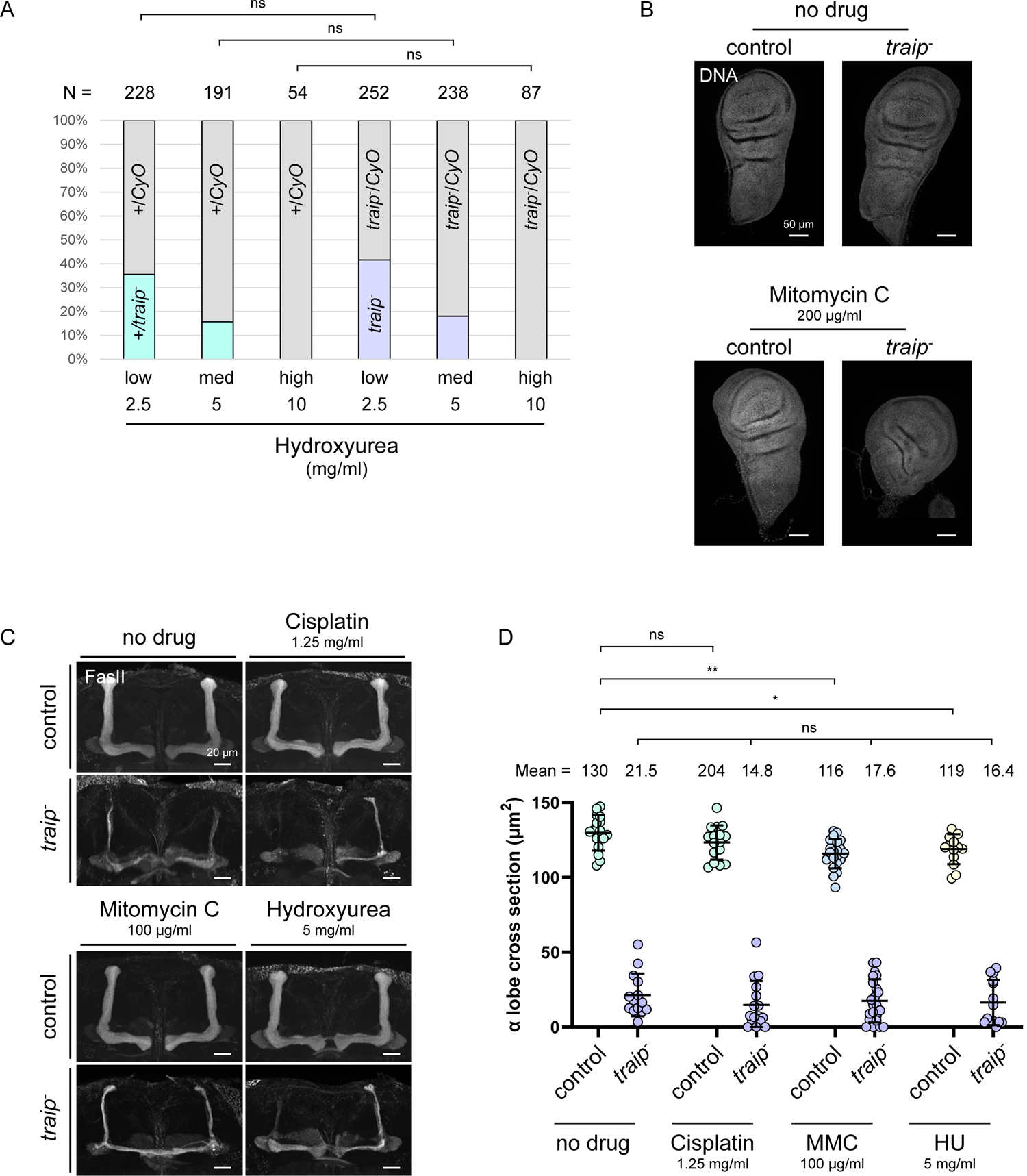
Traip is required for inter-strand crosslink repair, supporting figures. (A) Hydroxyurea treatment survival assay results. Both control and *traip^-^* crosses produced offspring that were increasingly sensitive to higher doses of hydroxyurea. Chi-squared tests were used for significance. (B) Wing discs from control and *traip^-^* 3^rd^ instar larvae treated with either no drug or high doses of Mitomycin C and stained for DAPI (gray). *traip^-^* discs treated with Mitomycin C are severely reduced in size compared to controls. (C) MBs from control and *traip^-^* treated with no drug or medium doses of Cisplatin, Mitomycin C, or Hydroxyurea, stained with FasII. Scale = 20 μm. (D) α lobe cross-section measurements of control and *traip^-^* treated with no drug or medium doses of Cisplatin, Mitomycin C (MMC), or Hydroxyurea (HU). Ordinary one-way ANOVA with Tukey’s test was used for significance. ns = not significant, ** p = 0.0013, * p = 0.0357. N ≥ 14 MBs.

**Figure S5.**
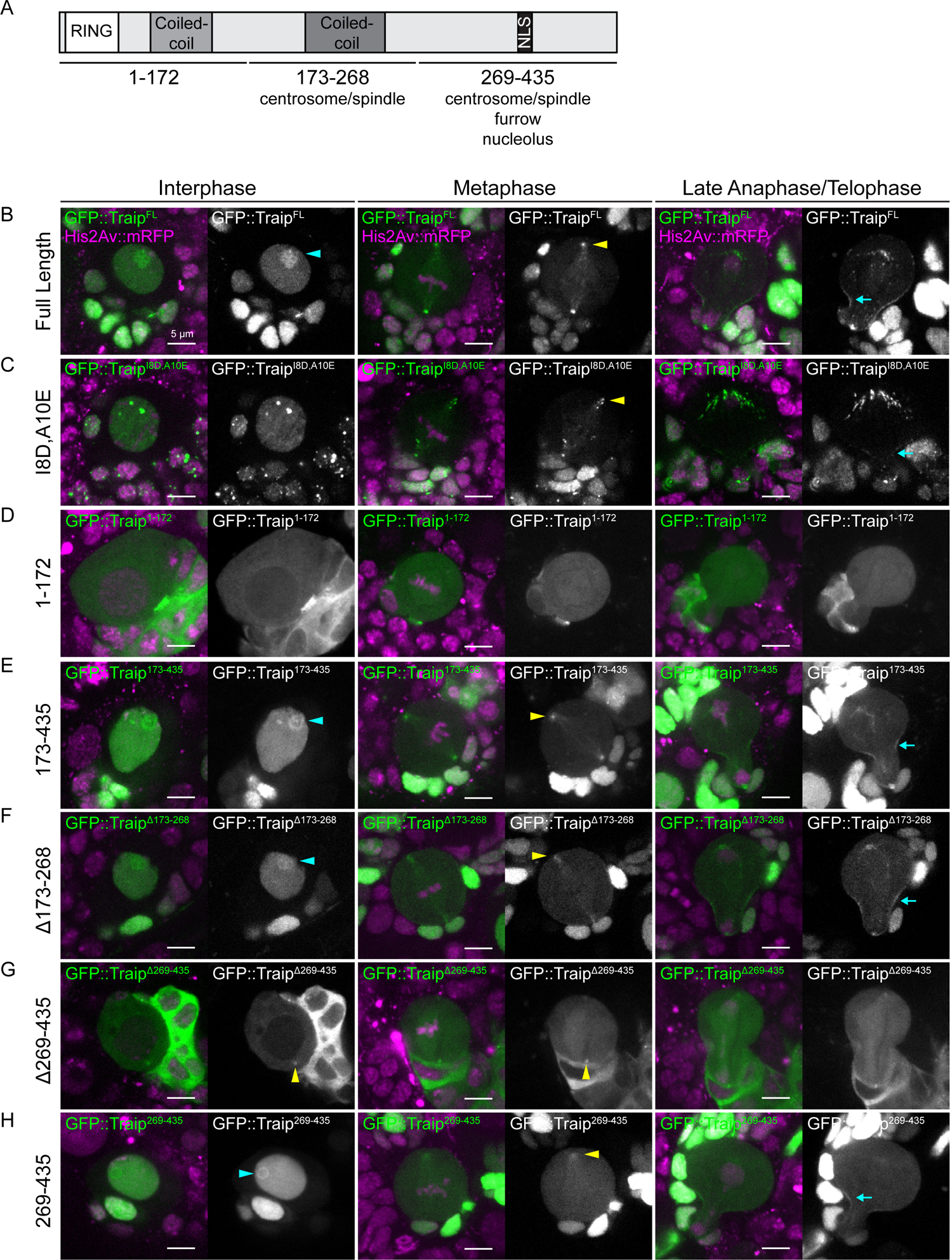
Traip localization depends on distinct domains. (A) Schematic of major Traip protein features, transgene fragment breakpoints, and localization domains. (B-H) Localization of GFP::Traip variant transgenes (green) expressed via *wor-GAL4* with His2Av::mRFP (magenta) in CB-NBs during interphase, metaphase, and late anaphase/telophase. Yellow arrowhead denotes centrosome localization, cyan arrow denotes cytokinetic furrow localization, and cyan arrowhead denotes nucleolar localization. GFP::Traip variants include: (B) Full Length, reproduced from Figure 7D; (C) RING domain mutant I8D, A10E; (D) RING domain and first coiled coil 1-172; (E) second coiled coil and C-terminal domain 173-435; (F) a deletion of the second coiled coil Δ173-268; (G) deletion of the C-terminal domain Δ269-435; and (H) the C-terminal domain alone 269-435. Scale bar = 5 μm.

**Figure S6.**
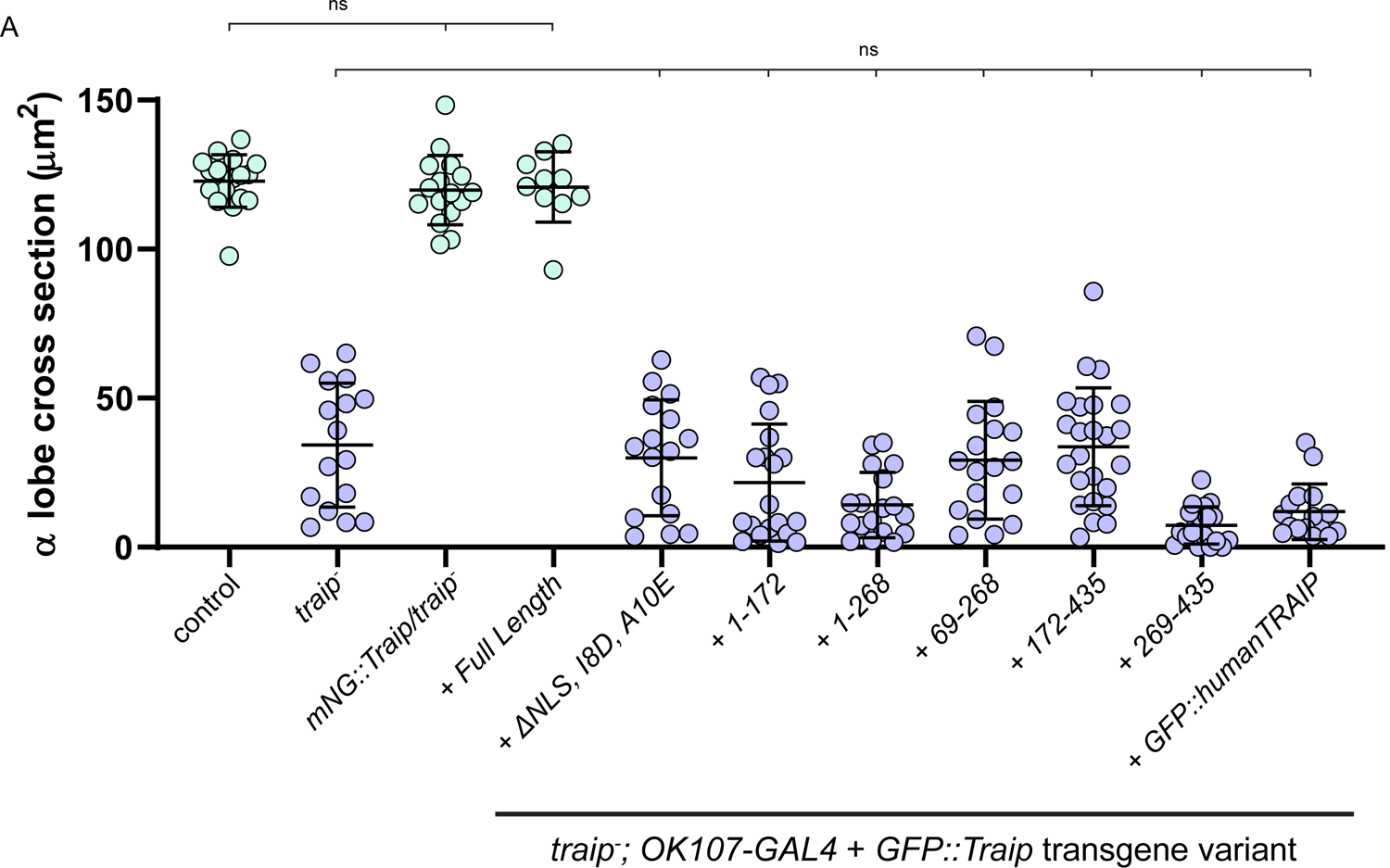
Transgene rescue negative results. (A) α lobe cross-section measurements of control, *traip^-^*, and *traip^-^* + GFP::Traip variant transgenes expressed via *OK107-GAL4*. Transgenes included *mNG::Traip^CRISPR^*, Full Length, ΔNLS + I8D,A10E, 1-172, 1-268, 69-268, 172-435, 269-435, and humanTRAIP. Ordinary one-way ANOVA with Tukey’s test was used for significance. ns = not significant. N ≥ 10 MBs.

**Figure S7.**
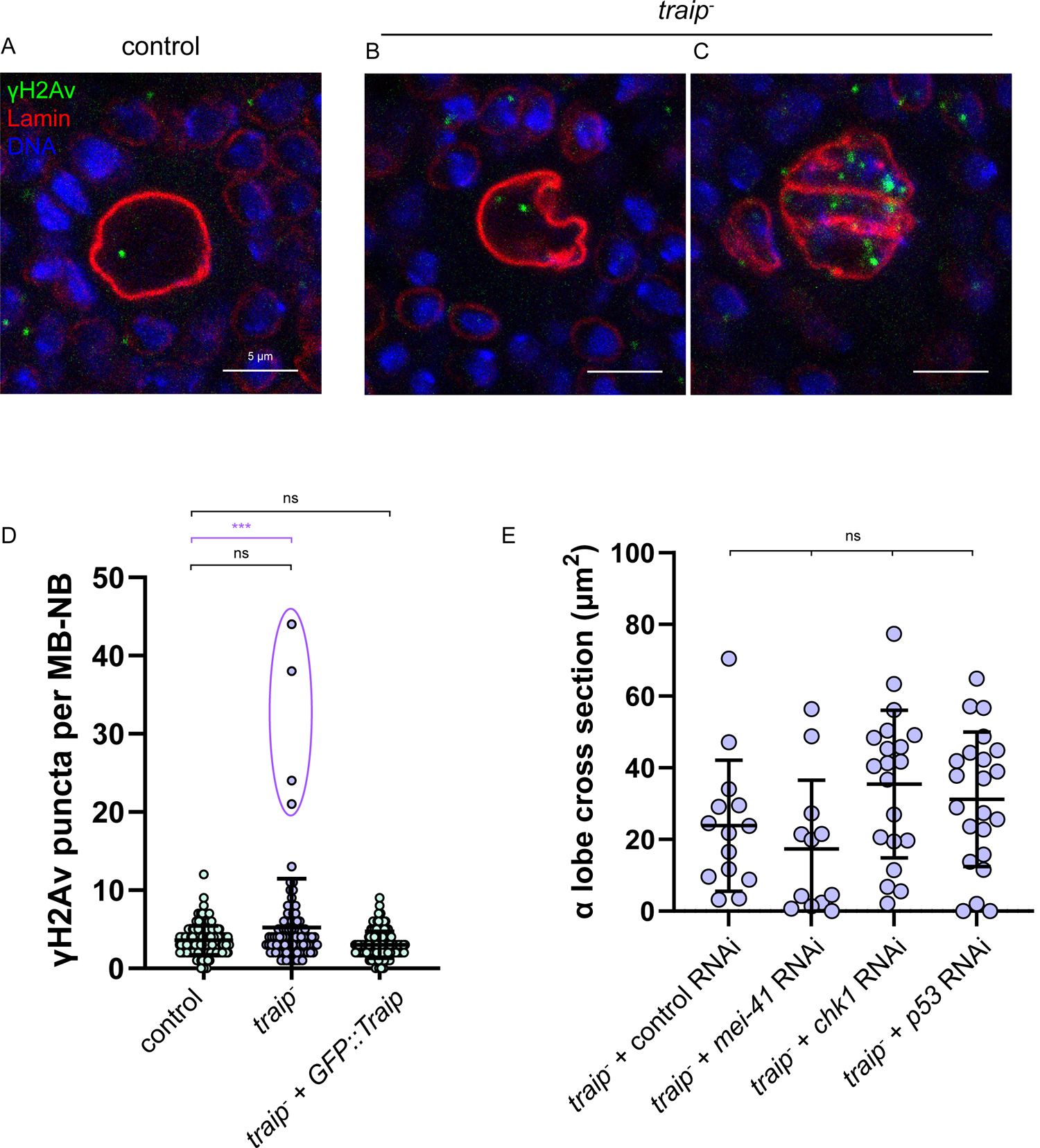
Traip does not function in interphase DNA damage repair. (A-C) Control and *traip^-^* 24 hours APF MB-NBs stained for γH2Av (green), Lamin (red), and DAPI (blue). Most control (A) and *traip^-^* (B) MB-NBs have few γH2Av puncta. Rare, likely apoptotic *traip^-^* MB-NBs have extremely elevated γH2Av puncta (C). (D) γH2Av puncta counts per MB-NB for control, *traip^-^*, and *traip^-^* + *GFP::Traip*. If *traip^-^* MB-NBs with elevated γH2Av puncta were included (purple outline), then *traip^-^* γH2Av puncta were significantly higher than controls (purple significance bar). If these *traip^-^* MB-NBs were excluded, then *traip^-^* γH2Av puncta were no longer elevated (black significance bar). Ordinary one-way ANOVA with test was used for significance. ns = not significant, *** p = 0.0004, **** p < 0.0001. N ≥ 100 MB-NBs. (E) α lobe cross-section measurements of *traip^-^* with control, *mei-41*, *chk1*, or *p53* RNAi expressed via *OK107-GAL4*. Two-tailed t-test was used for significance. ns = not significant. N ≥ 12 MBs.

